# A STING-CASM-GABARAP Pathway Activates LRRK2 at Lysosomes

**DOI:** 10.1101/2023.10.31.564602

**Authors:** Amanda Bentley-DeSousa, Agnes Roczniak-Ferguson, Shawn M. Ferguson

## Abstract

Mutations that increase LRRK2 kinase activity have been linked to Parkinson’s disease and Crohn’s disease. LRRK2 is also activated by lysosome damage. However, the endogenous cellular mechanisms that control LRRK2 kinase activity are not well understood. In this study, we identify signaling through Stimulator of Interferon Genes (STING) as an activator of LRRK2 via the Conjugation of ATG8 to Single Membranes (CASM) pathway. We furthermore establish that multiple chemical stimuli that perturb lysosomal homeostasis also converge on CASM to activate LRRK2. Although CASM results in the lipidation of multiple ATG8 protein family members, we establish that LRRK2 lysosome recruitment and kinase activation is highly dependent on interactions with the GABARAP member of this family. Collectively these results define a pathway that integrates multiple stimuli at lysosomes to control the kinase activity of LRRK2. Aberrant activation of LRRK2 via this pathway may be of relevance in both Parkinson’s and Crohn’s diseases.

## Introduction

Leucine rich repeat kinase 2 (LRRK2) has been linked by human genetics to Parkinson’s disease, Crohn’s disease and leprosy (Hui et al., 2018; Kalogeropulou et al., 2022; Kluss et al., 2019; Paisan-Ruiz et al., 2004; Rocha et al., 2022; Taylor and Alessi, 2020; Wallings and Tansey, 2019; Zhang et al., 2009; Zhu et al., 2023; Zimprich et al., 2004). LRRK2 mutations cause dominantly inherited familial Parkinson’s disease and common variants contribute to sporadic Parkinson’s disease risk (Taymans et al., 2023). It is also well established that multiple familial Parkinson’s disease LRRK2 missense mutations result in an increase in LRRK2 kinase activity (Kalogeropulou et al., 2022). Conversely, LRRK2 variants associated with decreased risk for Parkinson’s and Crohn’s disease exhibit reduced kinase activity (Wang et al., 2021). Although mutations that increase LRRK2 kinase activity are only found in a subset of Parkinson’s disease patients, the strong link between LRRK2 kinase activity and Parkinson’s disease suggests that other factors that modulate LRRK2 kinase activity could have disease relevant consequences.

Recent studies have converged on endosomes and lysosomes as key intracellular sites of LRRK2 activation. LRRK2 is recruited to lysosomes in response to chemical treatments that impair lysosome function and/or that cause lysosome membrane damage, and this is accompanied by an increase in the kinase activity of LRRK2 (Bonet-Ponce et al., 2020; Eguchi et al., 2018; Herbst et al., 2020; Kalogeropulou et al., 2020; Kluss et al., 2022; Radulovic and Stenmark, 2020). LRRK2 is also activated when pathogens such as *Mycobacterium tuberculosis*, *Listeria monocytogenes* or *Candida albicans* rupture lysosomes (Herbst et al., 2020). The dynamic recruitment of LRRK2 to perturbed endo-lysosomal membranes requires mechanisms whereby endo-lysosome status is sensed and communicated to LRRK2. Understanding the basis for this regulation has implications for understanding both the fundamental cellular functions of LRRK2 and potentially for how LRRK2 is aberrantly activated in disease contexts.

Severely damaged lysosomes are cleared by an autophagic process known as lysophagy wherein they are engulfed and delivered to healthy lysosomes for disposal (Maejima et al., 2013). However, cells can also detect and repair lysosome damage before it reaches the point where lysophagy is required (Bohannon and Hanson, 2020; Tan and Finkel, 2022). LRRK2 activation at damaged lysosomes has been proposed to promote the restoration of lysosome integrity by phosphorylating specific Rab GTPases that in turn recruit effectors involved in membrane repair (Bonet-Ponce et al., 2020; Herbst et al., 2020; Steger et al., 2016).

Rab GTPases also act upstream of LRRK2 to promote its membrane recruitment and activation (Dhekne et al., 2023; Eguchi et al., 2018; Gomez et al., 2019; Lian et al., 2023; Pfeffer, 2022; Purlyte et al., 2018; Unapanta et al., 2023; Vides et al., 2022; Wang et al., 2023a). However, although Rab GTPases promote LRRK2 kinase activity in specific contexts, recent knockout mouse studies revealed the persistence of partial or even full LRRK2 kinase activity in multiple tissues even after depletion of key Rab proteins implicated in LRRK2 activation (Dhekne et al., 2023; Kalogeropulou et al., 2020). These results suggest the existence of additional factors that contribute to LRRK2 activation.

Several Parkinson’s disease risk genes, including PINK1, Parkin, and VPS13C have been linked to the Stimulator of Interferon Genes (STING) signaling pathway that functions as part of an innate immune response downstream of the sensing of cytoplasmic DNA by cyclic GMP-AMP synthase (cGAS)(Diner et al., 2013; Hancock-Cerutti et al., 2022; Moehlman et al., 2023; Sliter et al., 2018; Wu et al., 2013). Motivated by this convergence of Parkinson’s disease genes and the fact that activated STING traffics through cellular compartments (Golgi, endosomes and lysosomes) where LRRK2 has been proposed to function, we investigated the relationship between LRRK2 and STING signaling (Erb and Moore, 2020). This led to the discovery that STING activates LRRK2 through a process known as V-ATPase–ATG16L1—induced LC3B lipidation (VAIL) or more generally as Conjugation of ATG8 to Single Membranes (CASM) (Durgan and Florey, 2022; Figueras-Novoa et al., 2024; Fischer et al., 2020). Although CASM broadly defines a process that results in the lipidation of multiple ATG8 family members at the surface of damaged or stressed organelles, we found that LRRK2 activation is dependent on an interaction with the GABARAP at lysosomes. This GABARAP dependent process of LRRK2 activation is required for LRRK2 activation in response to diverse stimuli that perturb lysosome membranes.

## Results

### STING is an endogenous activator of LRRK2

Given previously reported links between cGAS-STING dysregulation and Parkinson’s disease-associated genes, we tested whether STING controls LRRK2 activity. We chose the RAW 264.7 mouse macrophage cell line for these experiments based on their robust endogenous expression of both STING and LRRK2 combined with their experimental tractability. Following treatment with STING agonists [2′3′-Cyclic GMP-AMP (cGAMP) and DMXAA (5,6-Dimethylxanthenone-4-acetic Acid)], we observed the expected increases in both STING and TBK1 phosphorylation that reflect the established role for STING as an activator of TBK1(Fig. 1A-E)(Gui et al., 2019). We also measured levels of Rab10 phosphorylation on threonine 73 as a readout for changes in LRRK2 kinase activity (Kalogeropulou et al., 2022; Lis et al., 2018; Steger et al., 2016). Activation of STING by either agonist led to an increase in phosphorylation of Rab10 and the timing of DMXAA-dependent LRRK2 activation also paralleled STING activation (Fig. 1B-E; Fig. S1A). This relationship between STING and LRRK2 was also observed in human induced pluripotent stem cell-derived macrophages where treatment with a synthetic STING agonist (diABZI) resulted in increased Rab10 phosphorylation (Fig. S1B). Returning to RAW 264.7 cells, we also assessed Rab12 phosphorylation and found that cGAMP and DMXAA treatments both increased the phosphorylation of this additional LRRK2 substrate (Fig. S1C, D). The absence of Rab10 phosphorylation following STING activation in LRRK2 KO cells demonstrated the essential role for LRRK2 in Rab phosphorylation downstream of STING activation (Fig. 1B-E). To further test whether these effects were directly due to LRRK2 kinase activity, we acutely treated cells with the LRRK2 inhibitor MLi-2 and found that it also abolished the STING-mediated increase in Rab10 phosphorylation (Fig. 1F). Treating cells with another LRRK2 inhibitor, GZD-824, yielded the same effect (Fig. S1E). We also observed a lack of STING-mediated Rab phosphorylation in cells with a knockin of the inactive LRRK2 T1348N mutation (Ito et al., 2007)(Fig. S1F). Finally, LRRK2 mediated Rab10 phosphorylation following DMXAA treatment occurred with a concentration dependence that paralleled STING activation (Fig. S1G). Altogether, these data demonstrate that STING initiates a response that results in the activation of LRRK2 kinase activity.

**Figure 1:**
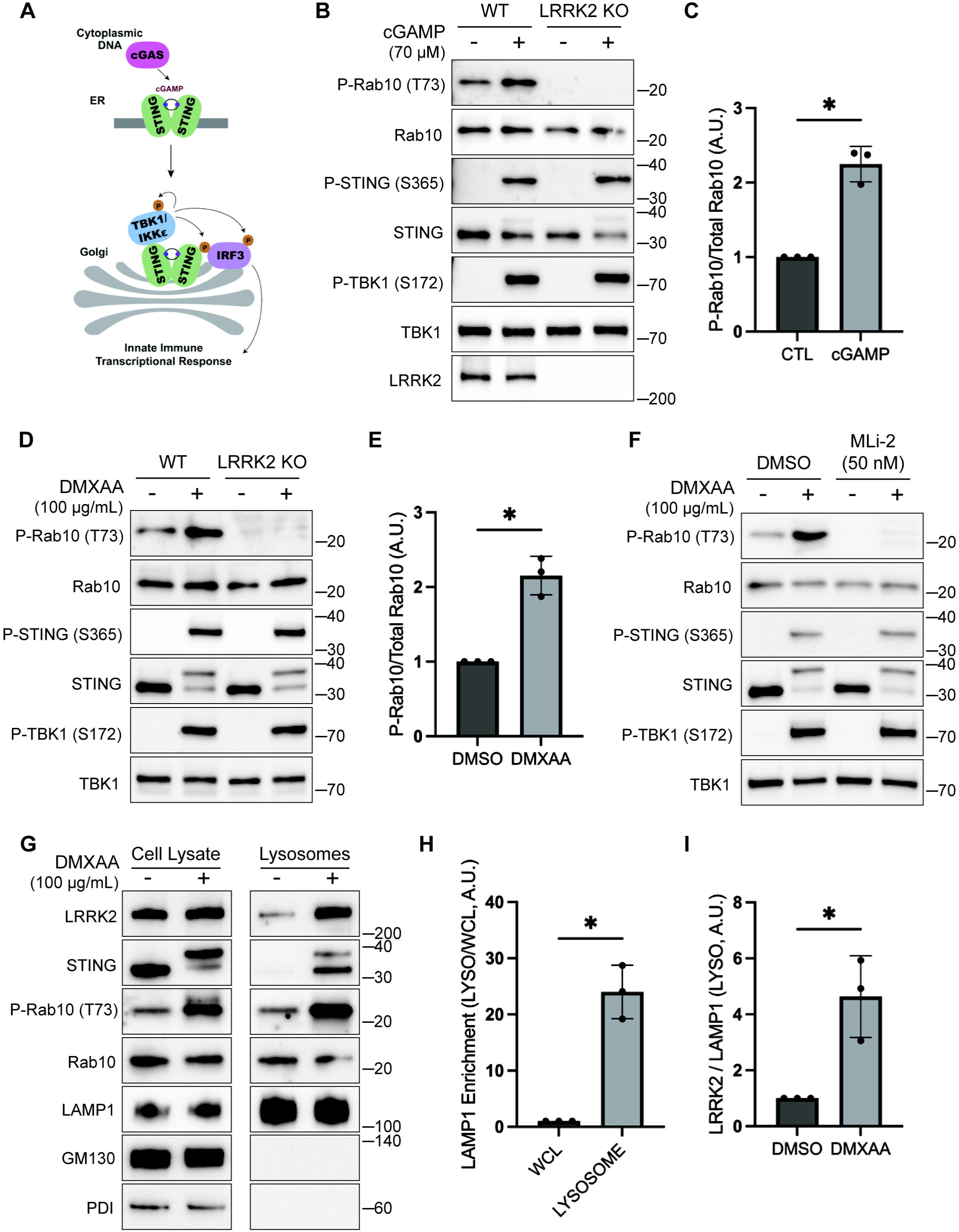
STING promotes LRRK2 kinase activity and lysosome recruitment. (A) Schematic representation of STING activation, trafficking and TBK1 activation. (B) Immunoblot analysis displaying the effects of a 2-hour treatment with cGAMP on the indicated proteins in wildtype and LRRK2 knockout cells. (C) Quantification of Rab10 phosphorylation in cGAMP-treated wildtype and LRRK2 knockout cells, with phospho-Rab10 levels normalized to total Rab10. A Two-Tailed Unpaired Welch’s t test was performed (P = 0.0118). (D) Immunoblot results demonstrating the impact of a 2-hour treatment with vehicle (DMSO) or DMXAA on the indicated proteins in cells of the specified genotypes. (E) Quantification of Rab10 phosphorylation in DMSO versus DMXAA-treated cells, with phospho-Rab10 levels normalized to total Rab10. A Two-Tailed Unpaired Welch’s t test was performed (P = 0.0163). (F) Immunoblot analysis depicting the effects of a 2-hour treatment with DMXAA on the indicated proteins in cells that were also treated with and without MLi-2. MLi-2 was added 1 hour before the addition of DMXAA and maintained during the DMXAA treatment. (G) Immunoblots of whole cell lysates and SPION-purified lysosomes from cells treated with DMSO or DMXAA for 2 hours. Results for the cell lysate and lysosome fractions were derived from the same membranes. Direct comparisons of organelle markers (LAMP1, GM130, PDI) are supported by the fact that these images reflect the same exposures and post-acquisition adjustments. (H) Quantification of LAMP1 enrichment in lysosome versus whole cell lysate (WCL) fractions to demonstrate effective lysosome enrichment. A Two-Tailed Unpaired Welch’s t test was performed (P = 0.0140). (I) Quantification of LRRK2 enrichment in the lysosome fractions (normalized to LAMP1 in the same fraction). A Two-Tailed Unpaired Welch’s t test was performed (P = 0.0496). Numbers at right of each blot refer to molecular weight (kDa). Error bars represent standard deviations. All experiments were performed in RAW 264.7 cells. Data presented in this figure is representative of results from a minimum of 3 independent experiments.

### STING promotes LRRK2 recruitment to lysosomes

Previous studies have shown that upon ligand binding, STING traffics from the endoplasmic reticulum to the Golgi and then to lysosomes where it is internalized and degraded (Balka et al., 2023; Chen et al., 2016; Gonugunta et al., 2017). Consistent with the known requirement for ER to Golgi trafficking as a first step in STING activation, we observed that disruption of such trafficking by treating cells with Brefeldin A prevented STING signaling, as was previously reported (Ishikawa et al., 2009) (Fig. S1H). We also found that Brefeldin A blocked LRRK2 activation downstream of STING (Fig. S1H). Meanwhile, LRRK2 can be activated at lysosomes in response to various lysosome damaging stimuli (Bonet-Ponce et al., 2020; Eguchi et al., 2018; Herbst et al., 2020; Kalogeropulou et al., 2020). To test the role of lysosomes as sites of STING-dependent LRRK2 activation, we next took advantage of an established method for isolation of lysosomes based on an endocytic pulse and chase of superparamagnetic iron oxide nanoparticles (SPIONs) followed by cell rupture and magnetic capture of lysosomes that contain the SPIONs (Amick et al., 2018; Amick et al., 2020; Hancock-Cerutti et al., 2022; Talaia et al., 2024). Comparison of lysosomes isolated from control versus DMXAA stimulated cells revealed that endogenous LRRK2 accumulated on lysosomes in response to STING activation (Fig. 1G-I). This was accompanied by phosphorylation of Rab10 at these lysosomes. Furthermore, we did not observe major changes in lysosome morphology in cells treated with DMXAA, unlike the changes in lysosome morphology observed with lysosome damaging agents (Fig. S2A, B).

### TBK1 and IKKε are not required for LRRK2 activation by STING

We next implemented a series of genetic perturbations to understand how STING signaling leads to LRRK2 activation. As expected, STING KO cells were defective in STING agonist-mediated LRRK2 activation and this was rescued following re-expression of STING (Fig. 2A). Following agonist binding, STING recruits and activates the closely related TBK1 and IKKε kinases (Balka et al., 2020; Chen et al., 2016)(Fig. 1A). Interestingly, TBK1 and IKKε were previously shown to phosphorylate LRRK2 which led us to speculate that this could be part of the pathway whereby STING activates LRRK2 (Dzamko et al., 2012). To test the requirement for TBK1 and IKKε in LRRK2 activation, we generated TBK1 KO, IKKε KO, and TBK1 + IKKε double KO RAW 264.7 cells and measured their ability to activate LRRK2 in response to STING agonist. In each of these KO lines, STING still triggered LRRK2-mediated Rab10 phosphorylation to a degree that was statistically indistinguishable from WT cells (Fig. 2B and C). Furthermore, we still observed LRRK2-mediated Rab10 phosphorylation after rescuing the STING KO cells with a truncated form of STING (amino acids 1-339) that cannot bind and activate TBK1/IKKε (Gui et al., 2019)(Fig. 2A). These results indicate that STING activates LRRK2 independent of signaling through TBK1 and IKKε kinases.

**Figure 2:**
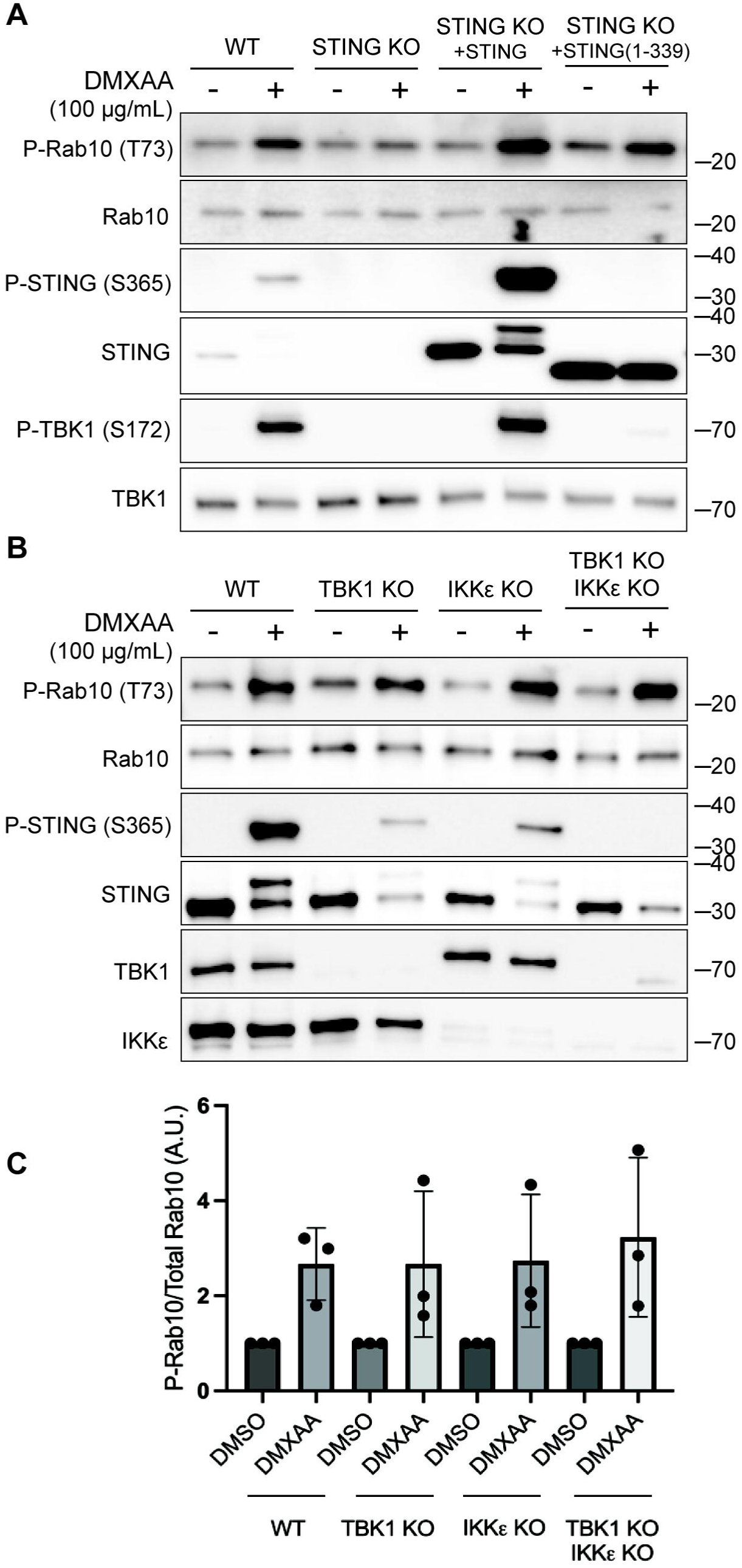
TBK1 and IKKE are not required for LRRK2 activation by STING. (A) Immunoblots revealing the effects of 2-hour treatments with DMSO or DMXAA on the indicated proteins in wildtype and STING knockout RAW 264.7 cells, along with STING knockout cells that were rescued by stable expression of full-length mouse STING or C-terminally truncated STING (amino acids 1-339). (B) Immunoblot results depicting the effects of 2-hour treatment with vehicle (DMSO) or DMXAA on the indicated proteins in wildtype, TBK1 knockout, IKKε knockout, and TBK1 + IKKε double knockout cells. (C) Quantification of Rab10 phosphorylation in DMSO versus DMXAA-treated cells of the indicated genotypes, with phospho-Rab10 levels normalized to total Rab10. A One-Way ANOVA was performed (P = 0.9476). Numbers at right of each blot refer to molecular weight (kDa). Error bars represent standard deviations. Data presented in this figure is representative of results from a minimum of 3 independent experiments.

### STING activates LRRK2 via the CASM pathway

In addition to TBK1-IKKε activation, STING also independently causes the lipidation of ATG8 family proteins in a non-autophagic process known as CASM (<u>C</u>onjugation of <u>A</u>TG8 to <u>S</u>ingle <u>M</u>embranes) and which has also been referred to by other names including VAIL (V-ATPase–ATG16L1—induced LC3B lipidation) (Durgan and Florey, 2022; Fischer et al., 2020; Gui et al., 2019; Hooper et al., 2022)(Fig. 3A). We observed that LRRK2 activation by STING ligands was also accompanied by both LC3B and GABARAP lipidation (Fig. 3B). ATG16L1 is a scaffold protein that is required for this lipidation reaction (Durgan and Florey, 2022; Fletcher et al., 2018; Hooper et al., 2022; Timimi et al., 2024). We therefore generated ATG16L1 KO RAW 264.7 cells, tested their response to STING activation, and found that LRRK2-mediated Rab phosphorylation was abolished (Fig. 3C). This observation led us to focus more deeply on a potential requirement for CASM in LRRK2 activation.

**Figure 3:**
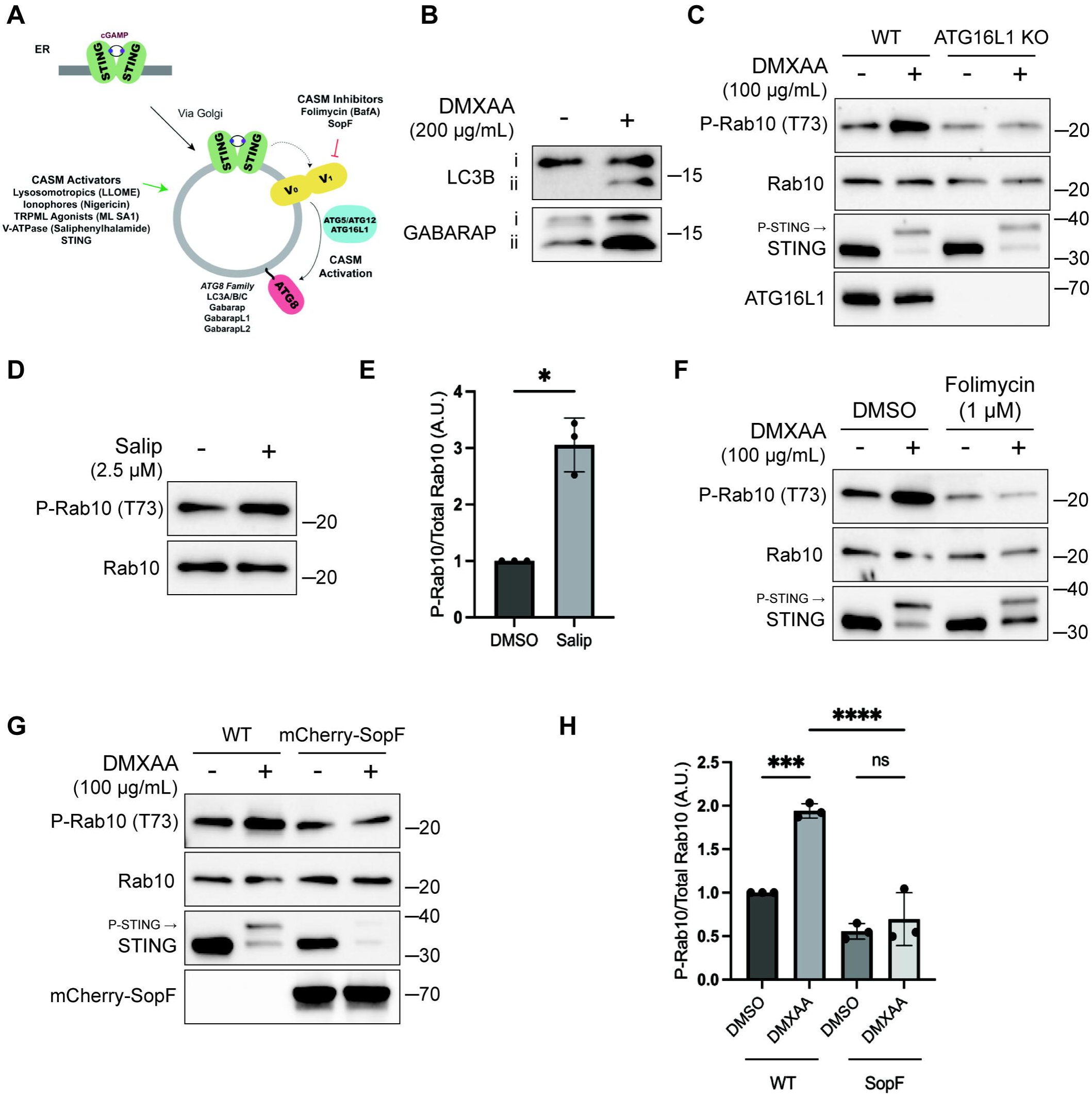
CASM is required for LRRK2 activation by STING. (A) Schematic diagram summarizing activators and inhibitors of the CASM pathway that were used in experiments in this figure. (B) Immunoblots illustrating the impact of DMXAA treatment on LC3B and GABARAP lipidation in RAW 264.7 cells (i = unlipidated, ii = lipidated). (C) Immunoblots displaying the effects of a 2-hour treatment with DMSO versus DMXAA on the indicated proteins in wildtype and ATG16L1 knockout cells. (D) Immunoblot results comparing control versus saliphenylhalimide (Salip) treated cells following a 2-hour treatment. (E) Quantification of the phospho-Rab10/total Rab10 ratio from panel D. A Two-Tailed Unpaired Welch’s t test was performed (P = 0.0175). (F) Immunoblots depicting the impact of Folimycin (added 1 hour before DMXAA and maintained throughout DMXAA incubation) on the response to a 2-hour treatment with DMSO versus DMXAA on the indicated proteins. (G) Immunoblots from wildtype RAW 264.7 cells versus cells that stably express mCherry-SopF after a 2-hour treatment with DMSO versus DMXAA, showing the impact on the indicated proteins in RAW 264.7 cells. (H) Quantification of the phospho-Rab/Rab10 ratios from panel G. A One-Way ANOVA with Sidak’s post-test was performed (WT DMSO vs WT DMXAA, P = 0.0006; WT DMXAA vs SopF DMXAA, P = <0.0001; SopF DMSO vs SopF DMXAA, P = 0.9023). Molecular weight markers (kDa) are indicated on the right side of each blot. Error bars represent standard deviations. All experiments were conducted in RAW 264.7 cells, and the data presented in this figure is representative of results obtained from a minimum of three independent experiments.

The V-ATPase plays a critical role in the CASM pathway and strategies have been defined that target the v-ATPase to activate or inhibit CASM (Durgan and Florey, 2022; Hooper et al., 2022; Timimi et al., 2024)(Fig. 3A). Saliphenylhalamide (Salip) leads to CASM by inhibiting the v-ATPase and stabilizing assembly of the V1 and V0 v-ATPase subunits (Hooper et al., 2022; Xie et al., 2004). Consistent with a role for CASM in activating LRRK2, Salip treatment increased LRRK2-mediated Rab10 phosphorylation (Fig. 3D and E). In contrast, treatment of cells with Folimycin (also known as Concanamycin A), a v-ATPase inhibitor that binds to a different site than Salip and inhibits CASM (but not the induction of macroautophagy), blocked the activation of LRRK2 downstream of STING (Fig. 3F) (Hooper et al., 2022). We also generated a cell line that stably expresses the Salmonella effector protein SopF which inhibits CASM by ADP-ribosylating the v-ATPase V_0_ subunit (Fischer et al., 2020; Hooper et al., 2022; Xu et al., 2019) and found that this abolished STING-dependent LRRK2 activation (Fig. 3G and H). Consistent with the proposed mechanism of action, SopF also inhibited the STING-induced lipidation of ATG8 family members (Fig. S3A). To rule out the involvement of macroautophagy in STING-induced LRRK2 activation, we knocked out FIP200, a protein that is essential for the ATG8 lipidation associated with macroautophagy but not for CASM and observed that STING-dependent LRRK2 activation still occurred in the absence of FIP200 (Fig. S3B and C) (Durgan and Florey, 2022; Figueras-Novoa et al., 2024; Fischer et al., 2020). Furthermore, STING also still activated LRRK2 kinase activity after treatment of cells with SAR405, a VPS34 inhibitor that blocks macroautophagy (Fig. S3D). These tests distinguished between ATG8 family lipidation associated with macroautophagy versus CASM and collectively support an essential role for CASM in mediating LRRK2 activation in response to multiple stimuli.

### Multiple chemicals activate LRRK2 via CASM

Multiple chemical stimuli that perturb lysosomes have separately been shown to activate LRRK2 and to activate CASM. This includes LLOME, a chemical that damages lysosome membranes following its processing by cathepsin C (Bonet-Ponce et al., 2020; Durgan and Florey, 2022; Kalogeropulou et al., 2020; Thiele and Lipsky, 1990). We confirmed that LLOME activates LRRK2 and established that LLOME also strongly triggers endogenous LRRK2 accumulation on lysosomes and activation in RAW 264.7 cells (Fig. 4A-C).

**Figure 4:**
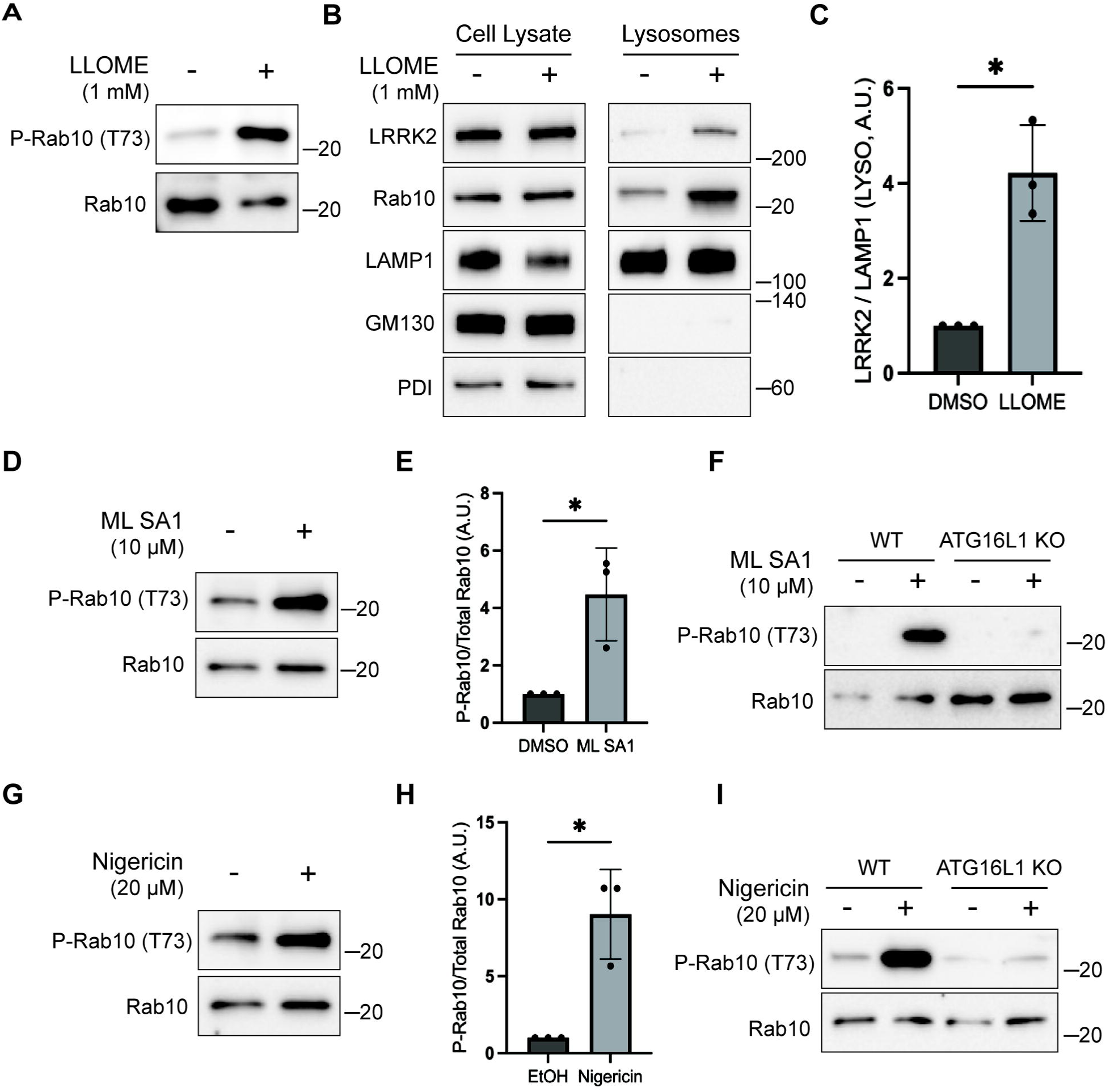
Multiple stimuli activate LRRK2 through CASM. (A) Immunoblots illustrating the impact of LLOME treatment (2 hours) on phospho-Rab10 and total Rab10 levels. (B) Immunoblots of cell lysates and SPION-purified lysosomes from cells treated with LLOME for 2 hours. Results for the cell lysate and lysosome fractions were derived from the same membranes. Direct comparisons of organelle markers (LAMP1, GM130, PDI) are supported by the fact that these images furthermore reflect the same exposures and post-acquisition adjustments. (C) Quantification of LRRK2 enrichment on lysosomes (normalized to LAMP1) from panel B. A Two-Tailed Unpaired Welch’s t test was performed (P = 0.0313). (D) Immunoblots demonstrating the effects of a 2-hour treatment with DMSO versus ML SA1 on Rab10 phosphorylation. (E) Quantification of phospho-Rab10/Rab10 ratios from panel D. A One-Tailed Unpaired Welch’s t test was performed (P = 0.0327). (F) Immunoblots showing the effects of a 2-hour treatment with ML SA1 on phospho-Rab10 and Rab10 in wildtype and ATG16L1 knockout cells. (G) Immunoblots displaying the effects of a 2-hour treatment with Nigericin on Rab10 phosphorylation. (H) Quantification of phospho-Rab10/Rab10 ratios from panel G. A Two-Tailed Unpaired Welch’s t test was performed (P = 0.0411). (I) Immunoblots illustrating the effects of a 2-hour treatment with Nigericin on phospho-Rab10 and Rab10 in wildtype and ATG16L1 knockout cells. Molecular weight markers (kDa) are indicated on the right side of each blot. Error bars represent standard deviations. All experiments were conducted in RAW 264.7 cells, and the data presented in this figure is representative of results obtained from a minimum of three independent experiments.

We next tested additional stimuli for their ability to induce both CASM and LRRK2 activation. TRPML1 is a lysosomal cation channel and a known activator of CASM but was not previously shown to activate LRRK2 (Durgan and Florey, 2022; Shen et al., 2012). We found that treatment with the TRPML1 agonist known as ML SA1 also activated LRRK2 and that this effect required ATG16L1 (Fig. 4D-F). Nigericin, a proton-potassium ionophore that disrupts ion balances across cellular membranes, is also known to cause CASM and to activate LRRK2 via unknown mechanisms (Herbst et al., 2020; Hooper et al., 2022; Ito et al., 2016; Jacquin et al., 2017; Kalogeropulou et al., 2020). We confirmed this robust relationship between Nigericin and LRRK2 kinase activity (Fig. 4G, H). Importantly, the KO of ATG16L1 blocked the ability of Nigericin to activate LRRK2 (Fig. 4I). As a control, we confirmed that these chemicals that were previously shown to cause CASM resulted in lipidation of ATG8 family members under the conditions where we observed LRRK2 activation (Fig. S3E). Collectively, these results support a role for CASM in the activation of LRRK2 by STING as well as by multiple lysosome perturbing chemical stimuli that induce CASM.

### GABARAP is required for CASM-dependent LRRK2 activation

To define specific cellular machinery that is required for CASM-mediated LRRK2 activation, we performed a targeted siRNA screen focusing on the ATG8-related proteins that are lipidated as part of the CASM process. As expected, DMXAA-induced LRRK2 activity (measured by Rab10 phosphorylation) was reduced by the knockdown of LRRK2 and Rab10 itself, as well as by knockdown of Rab12, a recently identified regulator of LRRK2 activity (Dhekne et al., 2023; Wang et al., 2023a)(Fig. 5A). Consistent with a requirement for lipidation of ATG8 family members, depletion of ATG3 and ATG16L1 also reduced the phosphorylation of Rab10 by LRRK2 (Fig. 5A). Interestingly, out of the ATG8 family members annotated in the mouse genome (LC3A, LC3B, GABARAP, GABARAPL1, and GABARAPL2) only GABARAP was identified as critical for CASM-mediated LRRK2 activation (Fig. 5A). This requirement for GABARAP in LRRK2 activation was independently validated through assays in genome edited GABARAP KO cells where neither DMXAA nor ML SA1 nor Nigericin were able to activate LRRK2 (Fig. 5B-E). Although there has previously been a major focus on lipidation of LC3 in association with the processes described as CASM and VAIL, the activators that we tested also caused the lipidation of GABARAP (Fig. S3G) (Fischer et al., 2020; Hooper et al., 2022; Liu et al., 2023). As evidence that GABARAP KO cells do not have major defects in processes that are upstream of CASM, we observed that GABARAP KO cells maintained STING-mediated LC3B lipidation (Fig. S4A). Additionally, STING phosphorylation and the phosphorylation of IRF3, a downstream target of STING signaling, were preserved in GABARAP KO cells (Fig. S4B). Defects in LRRK2 activation in the GABARAP KO cells were rescued by stable expression of HA-tagged GABARAP (Fig. S4C, D).

**Figure 5:**
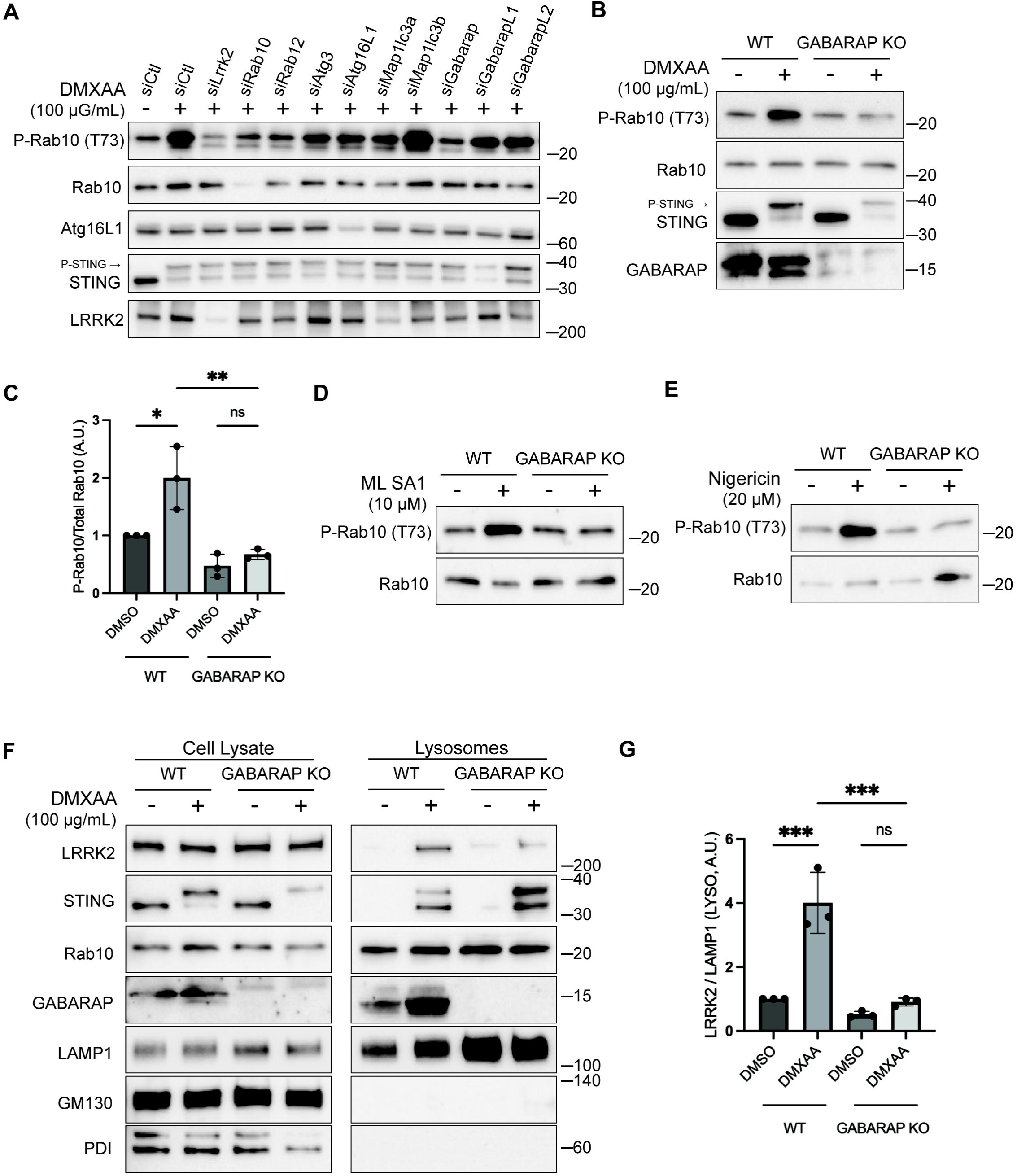
GABARAP is required for LRRK2 activation and lysosome recruitment. (A) Immunoblot results showing the impact of the indicated siRNAs on Rab10 phosphorylation and additional specified proteins in response to DMXAA. Experiments were performed 48 hours after siRNA transfections. (B) Immunoblots demonstrating the effects of a 2-hour treatment with DMXAA on the indicated proteins in wildtype and GABARAP knockout cells. (C) Quantification of the phospho-Rab10/Rab10 ratio in panel B. A One-Way ANOVA with Sidak’s post-test was performed (WT DMSO vs WT DMXAA, P = 0.0192; WT DMXAA vs GABARAP knockout DMXAA, P = 0.0034; GABARAP knockout DMSO vs GABARAP knockout DMXAA, P = 0.9650). (D) Immunoblots demonstrating the influence of GABARAP knockout on the phospho-Rab10/Rab10 ratio in response to ML SA1 (2-hour treatment). (E) Immunoblots revealing the effect of GABARAP knockout on the phospho-Rab10/Rab10 ratio in response to Nigericin (2-hour treatment). (F) Immunoblots of cell lysates and SPION-purified lysosomes from wildtype and GABARAP knockout cells treated with DMSO or DMXAA for 2 hours. Results for the cell lysate and lysosome fractions were derived from the same membranes. Direct comparisons of organelle markers (LAMP1, GM130, PDI) are supported by the fact that these images furthermore reflect the same exposures and post-acquisition adjustments. (G) Quantification of lysosomal LRRK2 abundance (LRRK2 normalized to LAMP1) from panel (F). A One-Way ANOVA with Sidak’s post-test was performed (WT DMSO vs WT DMXAA, P = 0.0004; WT DMXAA vs GABARAP knockout DMXAA, P = 0.0003; GABARAP knockout DMSO vs GABARAP knockout DMXAA, P = 0.9255). Molecular weight markers (kDa) are indicated on the right side of each blot. Error bars represent standard deviation. All experiments were conducted in RAW 264.7 cells, and the data presented in this figure is representative of results obtained from a minimum of three independent experiments, except for panel A where n=2.

Consistent with a role for GABARAP in recruiting LRRK2 to lysosomes as part of the activation process, analysis of purified lysosomes revealed that GABARAP is present on lysosomes following STING activation and is required for LRRK2 recruitment to lysosomes (Fig. 5F, G). GABARAP recruitment to lysosomes in response to CASM-inducing stimuli was also observed by immunofluorescence (Fig. S5A, B). Overall, our results demonstrate that GABARAP is critical for LRRK2 lysosomal recruitment and activation.

### GABARAP interacts with LRRK2 via distinct LIR motifs

Consistent with a potential direct role for GABARAP in promoting LRRK2 recruitment to lysosomes, Halo-tagged LRRK2 was detected in HA-GABARAP immunoprecipitations and this co-purification increased in response to STING activation (Fig. 6A). Figure S5C shows that the Halo-tagged LRRK2 is functional based on its ability to support both basal and DMXAA-stimulated Rab10 phosphorylation when stably expressed in LRRK2 KO cells. ATG8 family members often interact with other proteins via LIR/AIM (LC3-interacting region/ATG8-interacting motifs) and GABARAP interacting motifs (GIMs) (Rogov et al., 2017). Although there are no perfect matches for the GIM consensus sequence ([W/F]-[V/I]-X_2_-V) in human LRRK2, there were 42 matches to the LIR motif ([W/F/Y]-X_2_-[I/L/V])(Fig. 6C). As the large number of candidates that precluded systematic testing, we next used AlphaFold (v2.2.4 multimer model and subsequently the AlphaFold Server) to predict putative sites of interaction between LRRK2 and GABARAP (Abramson et al., 2024; Tunyasuvunakool et al., 2021). Queries for the interaction between full length human LRRK2 (Uniprot: Q5S007) and GABARAP Uniprot: O95166) yielded the identification of two candidate GABARAP interaction sites that we selected for functional validation. The first site of interaction that was centered on a consensus LIR motif (LIR#1: _109_WEVL_112_) within a loop extending from the ARM domain of LRRK2 and the LIR-ATG8 docking site on GABARAP (Fig. 6C and D)(Rogov et al., 2023). The second candidate LRRK2-GABARAP interaction was mediated by another LIR motif (LIR#2: _875_WTFI_878_) located at the base of the large loop that extends from the LRR region of LRRK2 (Fig. 6C and E). To test the functional significance of these predictions, we mutated the critical first and last amino acids of each candidate LIR motif to alanine to yield AEVA and ATFA for LIR1 and LIR2 respectively. Our subsequent experiments revealed that mutations to either of these motifs reduced the LRRK2-GABARAP interaction in response to STING activation (Fig. 7A, B). These LRRK2 LIR mutants were also unable to fully support STING-mediated LRRK2 activation (Fig. 7C, D). Finally, these LIR motifs were also critical for LRRK2 recruitment to lysosomes in response to STING activation (Fig. 7E, F). Consistent with a generalizable role these LRRK2-GABARAP interactions in the activation of LRRK2, mutations to LIR motifs 1 and 2 also prevented the activation of LRRK2 by LLOME (Fig. 7G, H).

**Figure 6:**
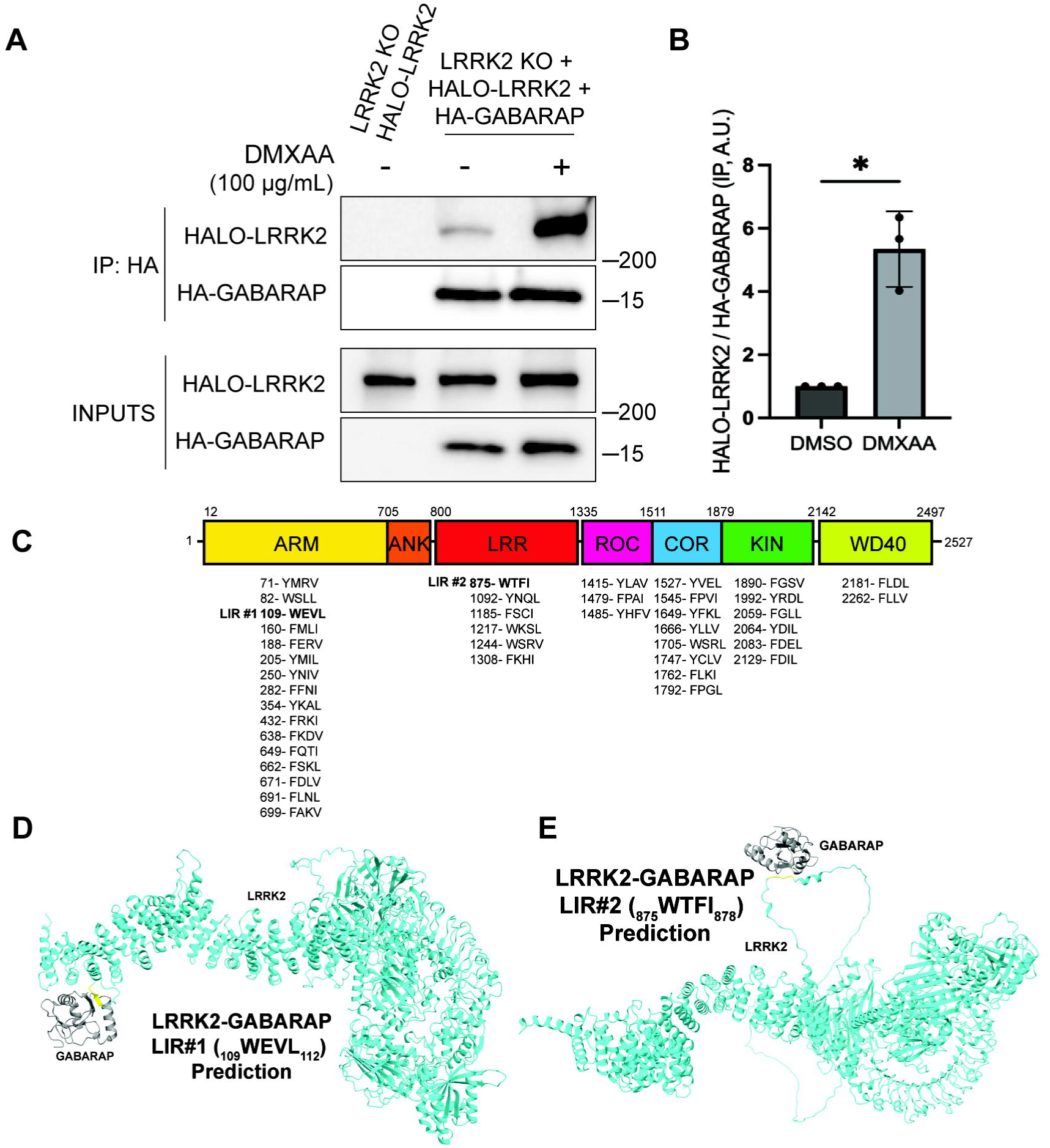
LRRK2 interacts with GABARAP via putative LIR motifs. (A) Immunoblots of the starting material (Inputs) and anti-HA immunoprecipitates from LRRK2 knockout cells that stably express Halo-tagged human LRRK2 +/-HA-tagged human GABARAP, treated for 2 hours with vehicle (DMSO) or DMXAA. (B) Quantification of Halo-LRRK2 abundance in HA-GABARAP immunoprecipitations +/-STING activation with DMXAA. A Two-Tailed Unpaired Welch’s t test was performed (P = 0.0243). (C) Schematic demonstrating the locations of all possible LIR motifs ([W/F/Y]-X_1_-X_2_-[I/L/V]) in LRRK2. (D and E) Structural predictions for interactions between LRRK2 (cyan) and GABARAP (grey) via LIR1 and LIR2 respectively. LIR motifs are highlighted in yellow. Molecular weight markers (kDa) are indicated on the right side of each blot. Error bars represent standard deviation. All experiments were conducted in RAW 264.7 cells, and the data presented in this figure is representative of results obtained from a minimum of three independent experiments.

**Figure 7:**
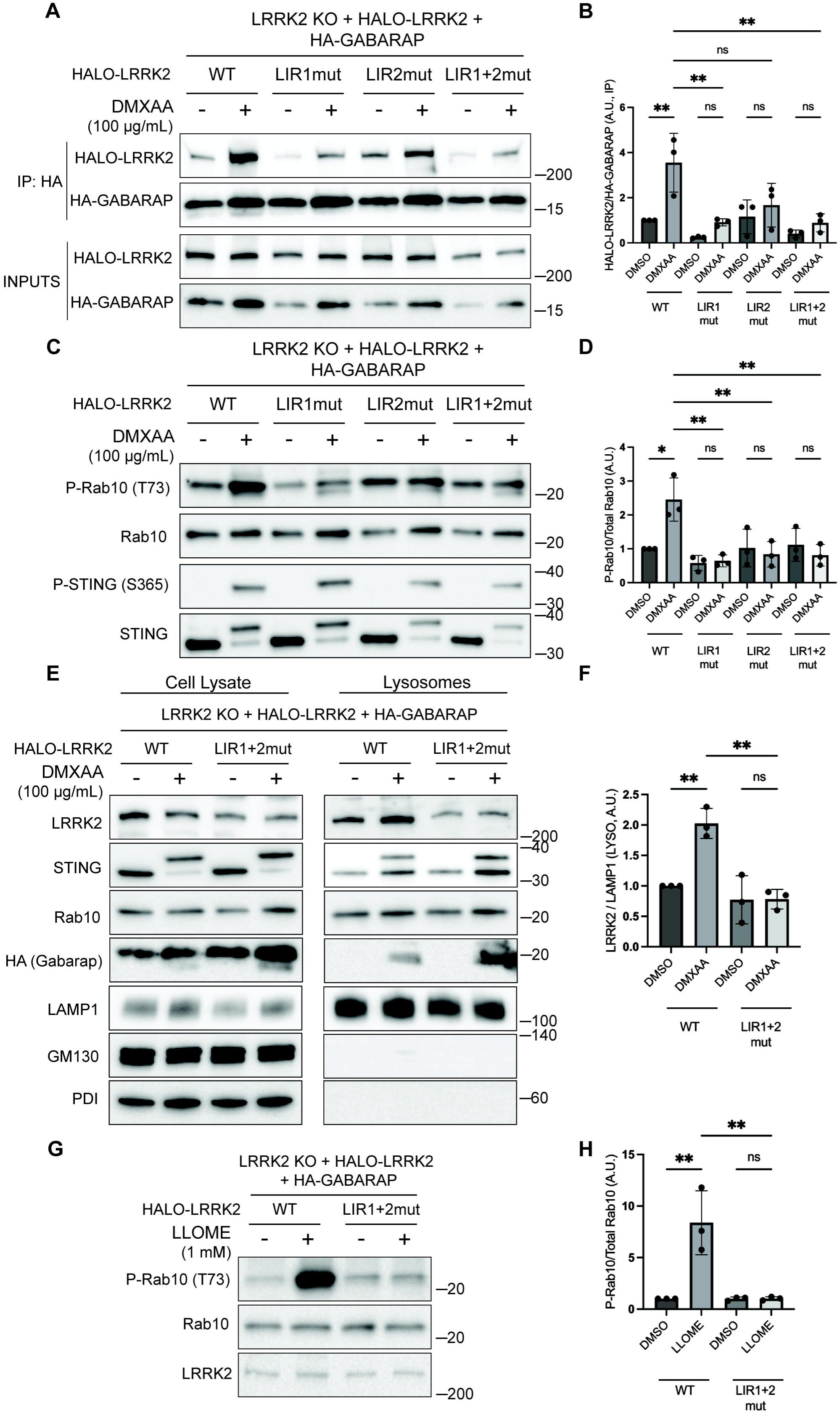
Functional validation of LRRK2 LIR motifs. (A) Immunoblots of inputs and anti-HA immunoprecipitates from LRRK2 knockout cells that stably express versions of Halo-tagged human LRRK2 (WT, LIR1 mutant, LIR2 mutant, and LIR1+2 mutant) and HA-tagged human GABARAP. LIR#1 mutation is WEVL to AEVA, LIR2 mutation is WTFI to ATFA, and LIR1+2 mutant combines LIR1 and LIR2 mutations. Cells were treated with DMSO or DMXAA for 2 hours. (B) Quantification of HALO-LRRK2 abundance in HA-GABARAP immunoprecipitations in (A). A One-Way ANOVA with Sidak’s post-test was performed (WT DMSO vs WT DMXAA, P = 0.0056; WT DMXAA vs LIR1mut DMXAA, P = 0.0040; WT DMXAA vs LIR2mut DMXAA, P = 0.0750; WT DMXAA vs LIR1+2mut DMXAA, P = 0.0037; LIR1mut DMSO vs LIR1mut DMXAA, P = 0.9993; LIR2mut DMSO vs LIR2mut DMXAA, P = >0.9999; LIR1+2mut DMSO vs LIR1+2mut DMXAA, P = >0.9999). (C) Immunoblots from LRRK2 knockout cells that stably express versions of Halo-tagged human LRRK2 (WT, LIR1 mutant, LIR2 mutant, and LIR1+2 mutant) and HA-tagged human GABARAP treated with DMSO or DMXAA (2 hours). (D) Quantification of phospho-Rab10 over total Rab10 in (C). A One-Way ANOVA with Sidak’s post-test was performed (WT DMSO vs WT DMXAA, P = 0.011; WT DMXAA vs LIR1mut DMXAA, P = 0.0012; WT DMXAA vs LIR2mut DMXAA, P = 0.0041; WT DMXAA vs LIR1+2mut DMXAA, P = 0.0034; LIR1mut DMSO vs LIR1mut DMXAA, P = >0.9999; LIR2mut DMSO vs LIR2mut DMXAA, P = >0.9999; LIR1+2mut DMSO vs LIR1+2mut DMXAA, P = >0.9999). (E) SPION-purified lysosomes from LRRK2 knockout cells stably expressing HALO-tagged human LRRK2 (WT and LIR1+2 mutant) +/-treatment with 100 µg/ml DMXAA (2 hours). Results for the cell lysate and lysosome fractions were derived from the same membranes. Direct comparisons of organelle markers (LAMP1, GM130, PDI) are supported by the fact that these images reflect the same exposures and post-acquisition adjustments. (F) Quantification of lysosomal LRRK2 abundance (LRRK2 normalized to LAMP1) from panel (E). A One-Way ANOVA with Sidak’s post-test was performed (WT DMSO vs WT DMXAA, P = 0.0056; WT DMXAA vs LIR1+2mut DMXAA, P = 0.0016; LIR1+2mut DMSO vs LIR1+2mut DMXAA, P = >0.9999). (G) Immunoblots from LRRK2 knockout cells that stably express versions of Halo-tagged human LRRK2 (WT or LIR1+2 mutant) and HA-tagged human GABARAP treated with DMSO or LLOME for 2 hours. (H) Quantification of phospho-Rab10 normalized to total Rab10 from panel (G). A One-Way ANOVA with Sidak’s post-test was performed (WT DMSO vs WT LLOME, P = 0.0024; WT LLOME vs LIR1+2mut LLOME, P = 0.0024; LIR1+2 DMSO vs LIR1+2 LLOME, P = >0.9999). Molecular weight markers (kDa) are indicated on the right side of each blot. Error bars represent standard deviation. All experiments were conducted in RAW 264.7 cells, and the data presented in this figure is representative of results obtained from a minimum of three independent experiments.

## Discussion

Collectively, our data defines a pathway for the activation of LRRK2 at lysosomes wherein STING and additional stimuli that perturb lysosome integrity converge on CASM-mediated GABARAP lipidation to recruit LRRK2 to lysosomes and this leads to an increase in LRRK2 kinase activity towards Rab substrates. Although there has been great progress in elucidating the CASM pathway, the physiological functions downstream of CASM have remained elusive (Cross et al., 2023; Durgan and Florey, 2022; Figueras-Novoa et al., 2024; Hooper et al., 2022; Jacquin et al., 2017; Timimi et al., 2024; Ulferts et al., 2021; Xu et al., 2019). Our identification of CASM-mediated, GABARAP-dependent, LRRK2 activation defines a new functional output of this pathway.

Given that GABARAP lipidation onto intracellular membranes occurs during both conventional macroautophagy and CASM, but only CASM activates LRRK2, an additional factor (or factors) likely contributes to the specificity of LRRK2 activation. Candidate contributors to a coincidence detection mechanism to ensure spatial control of LRRK2 activation at GABARAP-positive lysosomes include Rab GTPases that have also been demonstrated to mediate LRRK2 membrane recruitment and kinase activation (Pfeffer, 2022). These include Rab10, Rab12, Rab29, and Rab32 (Dhekne et al., 2023; Gustavsson et al., 2024; Hop et al., 2024; Purlyte et al., 2018; Vides et al., 2022; Wang et al., 2023a; Zhu et al., 2023). It remains to be determined whether GABARAP acts in parallel with any of these Rabs to ensure that LRRK2 is recruited to, and activated at, the correct intracellular membranes. It was also recently discovered that LRRK2 can assemble into helical polymers on membranes that are enriched in acidic lipid head groups (Wang et al., 2023b). Such LRRK2-membrane interactions could act together with GABARAP to specify sites of LRRK2 activation.

CASM can be subdivided into VAIL, which depends on the V-ATPase and ATG16L1 to recruit ATG5-ATG12 and a more recently described process that depends on sphingomyelin and TECPR1 for ATG5-ATG12 recruitment [known as sphingomyelin-TECPR1-induced LC3 lipidation (STIL)] (Boyle et al., 2023; Corkery et al., 2023; Figueras-Novoa et al., 2024; Kaur et al., 2023). Our data defines VAIL-dependent GABARAP lipidation as upstream of LRRK2 but does not rule out contributions from STIL. Within the broader CASM field, it remains to be determined whether VAIL and STIL recruit distinct effectors. To a large degree, progress on this front has been limited by progress in the identification of CASM effectors. With LRRK2 joining other proteins such as FLCN-FNIP and ATG2 as effectors of CASM, new opportunities are opening up to investigate the physiology controlled by CASM (Cross et al., 2023; Goodwin et al., 2021).

Our data supports a model wherein lipidated GABARAP directly interacts with LRRK2 to recruit it to lysosomal membranes (Fig. 8). The presence of two distinct LIR motifs that are both required for efficient LRRK2 activation downstream of CASM-inducing stimuli indicates that neither single LIR motif is sufficient to support LRRK2 lysosome recruitment and activation. This may reflect modest strength of each individual LIR motif interaction that is overcome by the avidity of dual GABARAP interaction sites. Future in vitro reconstitution and biophysical characterization of these interactions will help to test this hypothesis.

**Figure 8:**
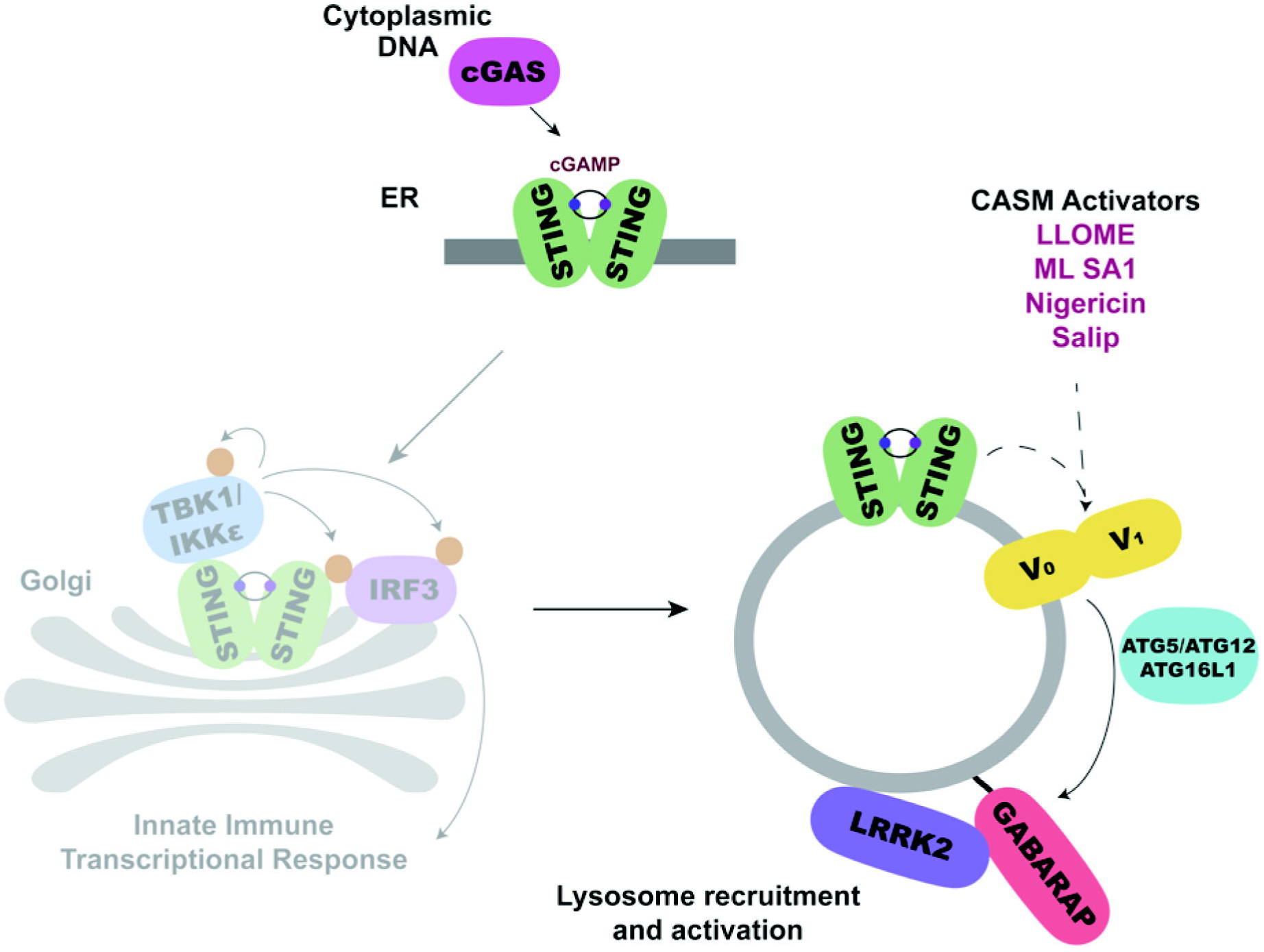
Schematic diagram summarizing how STING and additional CASM-inducing stimuli result in the lysosomal recruitment and activation of LRRK2 in a GABARAP-dependent manner.

The importance of LRRK2 kinase activity for Parkinson’s disease is well supported by the genetics of both sporadic and familial forms of the disease (Bandres-Ciga et al., 2020; Taylor and Alessi, 2020). The identification of the STING pathway as a robust endogenous activator of LRRK2 suggests that upstream stimuli that activate STING may also contribute to Parkinson’s disease risk. This includes cGAS-mediated synthesis of cGAMP in response to release of endogenous DNA from damaged mitochondria and disruptions to nuclear integrity as well as exogenous DNA from pathogen infections (Gulen et al., 2023; Lan et al., 2019; Motwani et al., 2019). Recent findings suggest that in the aging brain such effects may be of particular importance in microglia (Gulen et al., 2023). Additionally, failure of neuronal lysosomes to efficiently degrade nuclear and mitochondrial DNA can also result in STING activation following DNA leakage into the cytoplasm (Lan et al., 2014; Van Acker et al., 2023).

The role for CASM and GABARAP in LRRK2 activation provides a unifying mechanism that explains how diverse stimuli that perturb lysosome integrity and/or ion balance result in LRRK2 activation. In fact, the discoveries that STING itself is a proton channel suggests that it may activate CASM by disrupting endo-lysosomal pH (Liu et al., 2023; Xun et al., 2024). The identification of CASM as a convergence point for multiple stimuli that activate LRRK2 provides a foundation for further investigation of how lysosome dysfunction contributes to Parkinson’s disease. CASM-dependent LRRK2 activation may also be relevant for Crohn’s disease where human genetics has established roles for both ATG16L1 and LRRK2 and our data now places these genes into a common cellular pathway (Hampe et al., 2007; Van Limbergen et al., 2009).

In conclusion, we have defined a STING-CASM-GABARAP pathway for LRRK2 activation (Fig. 8). Our results more broadly explain how multiple additional lysosome perturbations converge to activate LRRK2. By establishing a site and machinery for LRRK2 activation, these results provide a new avenue for investigating the regulation of LRRK2 activity in the context of normal physiology as well as human diseases such as Parkinson’s disease, Crohn’s disease and microbial pathogenesis.

Note: While our manuscript was under revision, a separate study reported complementary results concerning the role of CASM in LRRK2 activation at lysosomes but without insights related to STING as an endogenous upstream activator or to the LRRK2-GABARAP interaction (Eguchi et al., 2024).

## Methods

### RAW 264.7 cell culture

RAW 264.7 cells were cultured in DMEM (Thermo Fisher Scientific, 11965-092) + 10% FBS (Thermo Fisher Scientific, 16140-071) + 1% penicillin/streptomycin supplement (Thermo Fisher Scientific, 15140122) at 37°C with 5% CO2. Cells were passaged using CellStripper (Corning, 356230). Details of cell lines are summarized in Supplemental Table 1. Sources of key reagents are listed in Supplemental Table 2.

### Human iPSC cell culture and macrophage differentiation

The A18945 human iPSC line (Thermo Fisher Scientific) was grown on Matrigel (Corning, 356230) coated dished in E8 media (Thermo Fisher Scientific, A15169-01) supplemented with E8 supplement (Thermo Fisher Scientific, A15171-01). Rock Inhibitor (Stemcell Technologies, 100-1044) was included overnight after passaging. A STEMdiff Hematopoietic Kit (Stemcell Technologies, 5310) was used for differentiation to hematopoietic progrenitor cells (HPCs). After 12 days, the HPCs were collected and grown in macrophage differentiation media (RPMI + 20% FBS + 100 ng/mL M-CSF (Peprotech, 30025) for 6 days (with a media change on day 15).

### Stable cell line generation

A piggybac transposase strategy was used for stable cell line generation. Briefly, 2.5 x 10^5^ cells were plated per well in a 6-well dish. The following day, cells were transfected using Lipofectamine 2000 (Invitrogen, 11668019) or FuGene HD (Promega, E2311), as per the manufacturer’s protocols, using a 1:2 ratio of Piggybac transposon plasmid:gene-of-interest plasmid (total 1 µ g DNA). After 48 hours, the media was changed. After 72 hours, puromycin (Gibco A11138-03) or blasticidin (Invivogen ant-bl) were used at concentrations of 3.5 µg/mL and 10 µg/mL, respectively. After 48-72 hours, media containing puromycin and blasticidin was washed out with fresh media. After cells had recovered from selection, single cells were plated into 96-well dishes to obtain clonal cell lines. Cell lines were confirmed via immunoblotting. Plasmid information is summarized in Supplemental Table 3.

Custom murine STING, human Halo-LRRK2, and human HA-GABARAP piggybac vectors were purchased from VectorBuilder. The C-terminal truncation of murine STING (to include only amino acids 1-339) was accomplished using NEB Q5 High-Fidelity 2X Master Mix (NEB M0492S) per the manufacturer’s protocol. Plasmids were sequenced to confirm successful mutagenesis. pmCherry-SopF was a gift from Leigh Knodler (Addgene plasmid # 135174; http://n2t.net/addgene:135174; RRID:Addgene_135174)(Lau et al., 2019). In order to create a stable cell line, the mCherry-SopF coding sequence was cloned into a pPB Piggybac plasmid (Vectorbuilder) using the NEB HIFI DNA Assembly Kit (NEB, E2621L). Plasmids will be made available via Addgene.

### Genome edited cell lines

STING KO, TBK1 KO, IKKe KO, TBK1 IKKc double KO, ATG16L1 KO, FIP200 KO and GABARAP KO cells were created using the Synthego CRISPR Gene Knockout V2 Mouse Kits for each respective target. Briefly, 2.5 x 10^5^ cells were plated into a 6-well dish. The following day, cells were transfected with ribonucleoprotein particles using Lipofectamine CRISPRiMAX (ThermoFisher Scientific, CMAX00003), gene specific sgRNAs and recombinant Cas9 (Synthego CRISPR Gene Knockout V2). After 48 hours, the media was changed. After 72 hours, single cells were plated into 96-well dishes to obtain clonal populations. After expansion of clonal populations, KO clones were identified by immunoblotting.

### Generation and use of superparamagnetic iron oxide nanoparticles (SPIONs)

SPIONS were generated based on an established protocol (Amick et al., 2018; Hancock-Cerutti et al., 2022; Rodriguez-Paris et al., 1993; Talaia et al., 2024; Wang et al., 2024)(https://www.protocols.io/view/synthesis-of-colloidal-dextran-conjugated-superpar-eq2lyn69pvx9/v1). Briefly, 10 mL of 1.2M FeCl_2_ (Sigma-Aldrich, 220299) and 10 mL of 1.8M FeCl_3_ (Sigma-Aldrich, 157740) were combined slowly by stirring. Then, 10 mL of 30% NH_4_OH (Sigma-Aldrich, 320145) was slowly added while stirring for 5 minutes. The resulting particles were then washed with 100 mL of water three times. The particles were resuspended in 80 mL of 0.3M HCl (J.T. Baker, 9535) and stirred for 30 minutes. Then, 4g of dextran (Sigma-Aldrich D1662) was added and stirred for 30 minutes. The particles were transferred into dialysis tubing and dialyzed with ddH_2_O for at least 2 days with multiple water changes. The particles were centrifuged at 26,900g for 30 minutes to remove large aggregates and stored at 4°C.

### SPION-mediated lysosome purification

An established SPION protocol for mouse macrophages was followed for lysosome purification with a few alterations (Rofe and Pryor, 2016). In short, a ∼90% confluent 15-cm dish was split into 8 10-cm dishes. The next day, 10 mL of fresh media containing 5% SPION particles supplemented with 5 mM HEPES (pH 7.4, Thermo Fisher Scientific, 15630-080) was added for 1 hour. The dishes were washed 2X with PBS. Fresh media was added to wash out the SPIONS for 2 hours. All following steps are on ice. Each dish was washed and scraped in PBS and centrifuged at 4°C for 5 minutes at 300 RCF. The cells were resuspended in 1 mL ice-cold HB buffer (5 mM Tris Base (American Bio, AB02000-05000), 250 mM Sucrose (Sigma-Aldrich, S0389), and 1 mM EGTA pH 7.4, Sigma-Aldrich, E4378) supplemented with inhibitors [cOmplete mini EDTA-free protease inhibitor (Roche, 11836170001) and PhosSTOP (Roche, 4906837001)] and transferred to a Dounce homogenizer. Cells were homogenized 50 times with the tight pestle and centrifuged at 4°C for 10 minutes at 800 RCF to obtain the cell lysate. At this step, a portion of the cell lysate was kept. LS columns (Miltenyi Biotec, 130042401) were washed once with 2.5 mL of HB buffer on a QuadroMACS separator (Miltenyi Biotec, 130-091-051). The remainder of the cell lysate was applied to LS columns. The flow-through was collected and reapplied on the column. The columns were washed with 3 mL of HB buffer, removed from the magnetic rack, and then lysosomes were eluted into ultracentrifuge tubes in 2.5 mL of HB buffer. Samples were ultracentrifuged at 55,000 RPM at 4°C for 10 minutes using a TLA-100.3 rotor in a Beckman-Coulter Ultracentrifuge Max Optima. The supernatant was removed, and the lysosome pellet was resuspended in ∼50 µl of HB buffer and prepared for immunoblotting.

### Immunoblotting

1 x 10^6^ cells were seeded per well in a 6-well dish. The following day, cells were washed 2 times with ice-cold 1X PBS [1.1 mM KH_2_PO_4_ (J.T. Baker, 3246-01), 155.2 mM NaCl (Sigma-Aldrich, 3624-05), and 3 mM Na_2_HPO_4_ (J.T. Baker, 3828-05)] and scraped in 50 µl of ice-cold lysis buffer [50 mM Tris Base (American Bio, AB02000-05000), 150 mM NaCl (Sigma-Aldrich, 3624-05), 1% Triton X-100 (Sigma-Aldrich, X100), 1 mM EDTA (Sigma-Aldrich, 03690) supplemented with cOmplete mini EDTA-free protease inhibitor (Roche, 11836170001) and PhosSTOP (Roche, 4906837001). Lysates were centrifuged at 14,000 RPM (4°C) for 8 minutes to remove insoluble material. Protein concentrations in the supernatants were measured using Coomassie Plus Protein Assay Reagent (ThermoFisher Scientific, 23236) as per manufacturer’s protocol. The lysate supernatants were mixed 1:1 with Laemmli Buffer [80 mM Tris-HCl pH 6.8 (American Bio, AB020000-05000 / HCl, Sigma-Aldrich, 3624-06), 25.3% Glycerol (American Bio, AB00751), 2.67% SDS (American Bioanalytical, AB01920-00500), and Bromphenol Blue (Sigma-Aldrich, B5525) supplemented with 6.187% fresh β-mercaptoethanol (Sigma-Aldrich, M3148)] and heated at 95°C for 3 minutes. Between 20-30 µg of protein was electrophoresed in 4-15% miniPROTEAN TGX Stain-Free pre-cast gels (BioRad, 4568084/4568085/4568086) using electrophoresis buffer [24.76 mM Tris Base (American Bio AB02000-05000), 191.87 mM Glycine (American Bio AB00730-05000), 10 mL 10% SDS (American Bio, AB01920-00500)]. Typically, 30 µg of protein was loaded for all protein lysates. For lysosome purifications, 20 µg of whole cell lysate was loaded, and 2 µg of lysosome sample was loaded. After electrophoresis, gels were transferred onto 0.45 µm pore nitrocellulose membrane (Thermo Fisher Scientific, 1620115) at 100V for 60 minutes in transfer buffer [24.76 mM Tris Base (American Bio AB02000-05000), 191.87 mM Glycine (American Bio AB00730-05000), 20% Methanol (Sigma-Aldrich, 179337-4l-pB)]. Total protein was visualized using BioRad TGX stain-free technology and/or Ponceau S prior to blocking. Membranes were blocked in 5% non-fat dry milk omniblock (American Bio, AB 10109-01000) in TBST [10 mM Tris Base (American Bio AB02000-05000), 150 mM NaCl (Sigma-Aldrich, 3624-05), 0.1% Tween 20 (Sigma-Aldrich, P7949)]. Antibodies were added in 5% BSA (Sigma-Aldrich, A9647) in TBST overnight (4°C) at the indicated concentrations (supplemental data Table 5). Membranes were washed 2 X 10 minutes with TBST. Secondary was added in TBST or 5% non-fat dry milk Omniblock in TBST (for phospho-specific antibody) at the indicated concentrations for 1 hour at room temperature. Membranes were washed 3 X 10 minutes with TBST. Membranes were subjected to chemiluminescence [SuperSignal West Pico PLUS Chemiluminescence Substrate (Thermo Fisher Scientific, 34580) or SuperSignal West Femto Maximum Sensitivity Substrate (Thermo Fisher Scientific, 34095)] and imaged using a Biorad Chemidoc MP imaging station. A summary of all antibodies used can be found in Table 5 in the supplemental data.

### Immunoprecipitation

For cell lysis, the immunoblotting protocol was followed with few alterations. One 80% confluent 15-cm plate was used per sample. RAW 264.7 LRRK2 KO cells rescued with stably expressed Halo-human LRRK2 (but no HA-GABARAP) were used as a negative control for non-specific LRRK2 interactions with the anti-HA beads. RAW 264.7 LRRK2 KO cells rescued with Halo-LRRK2 (WT as well as mutants of interest) and stably expressing HA-tagged human GABARAP were used to test LRRK2-GABARAP interaction. Cells were washed 2X with PBS, scraped in ice-cold lysis buffer, and centrifuged at 14,000 RPM (4°C) for 8 minutes. Protein concentrations were measured using Coomassie Plus Protein Assay Reagent (ThermoFisher Scientific, 23236) as per manufacturer’s protocol. Lysates were then immunoprecipitated using a mix of 15 µl anti-HA beads (Thermo Fisher Scientific, 88837) that were pre-washed 3X with lysis buffer. The same amount of protein was used for each sample. Where needed, samples were supplemented with lysis buffer to maintain the same protein concentration and volume. Lysates were incubated with beads rotating end-over-end for 1 hour at 4°C. Beads were washed 3X with 0.1% TBST and 1X with mqH20, as per the manufacturer’s protocol. Proteins were eluted by incubating beads with Laemmli Buffer and boiling at 95°C for 3 minutes. The sample was then transferred to another microcentrifuge tube and supplemented with 6.187% fresh β-mercaptoethanol (Sigma-Aldrich, M3148). Samples were then subjected to electrophoresis, immunoblotting, and chemiluminescence as previously described above.

### siRNA

siRNA-mediated knockdowns of target gene expression were accomplished using Horizon Biosciences siGENOME pooled siRNAs. Briefly, 2.5 x 10^5^ RAW 264.7 cells were plated per well in a 6-well dish. The following day, cells were transfected using Lipofectamine RNAiMAX (Invitrogen, 2448190) as per the manufacturer’s protocols using 100 nM siRNA pool. After 48 hours, cells were treated with drugs or lysed and subjected to immunoblotting. siRNAs used in this study are summarized in Supplemental Table 6.

### Immunofluorescence and imaging

2 x 10^5^ cells were plated on poly-D-lysine coated coverslips (12 mm, Carolina Biological Supplies, 633029). After treatment, cells were fixed in a 4% paraformaldehyde (Electron Microscopy Sciences, 19202)/sodium phosphate buffer (pH 7.3 / Buffer: 153.56 mM Sodium phosphate, dibasic, anhydrous, J.T. Baker 3828, 53.63 mM Sodium dihydrogen phosphate monohydrate, J.T. Baker 3818) for 30 minutes at room temperature. Cells were washed 3 times for 5 minutes each with PBS. Cells were blocked and permeabilized in 3% BSA in PBS supplemented with 0.1% Saponin (Sigma-Aldrich, S4521) for figure S2 or with 0.1% Triton for figures S5A and S5B (Sigma-Aldrich, S4521). Primary antibody was added overnight at 4°C. Cells were washed 3 times for 5 minutes each with PBS. Secondary antibody was added for 1 hour at room temperature in the dark. Cells were washed 3 times for 5 minutes each with PBS. Coverslips were mounted onto microscope slides (Thermo Fisher Scientific, 12-550-143) with Prolong Gold mounting media (Thermo Fisher Scientific, P36935) and stored at 4°C. For imaging, a Nikon Ti2-E inverted microscope with Spinning Disk Super Resolution by Optical Pixel Reassignment Microscope (Yokogawa CSU-W1 SoRa, Nikon) was used with a 60X SR Plan Apo IR objective with oil. Images were processed in FIJI.

### AlphaFold structural predictions

Using AlphaFold v2.2.4 multimer model as well as the AlphaFold Server, we modeled how full-length human LRRK2 would interact with human GABARAP. The predicted models were displayed and analyzed using UCSF ChimeraX-1.7.1, developed by the Resource for Biocomputing, Visualization, and Informatics at the University of California, San Francisco. ChatGPT-4o was used to generate a Python script that identified motifs in LRRK2 that conform to the [W/F/Y]-X_1_-X_2_-[I/L/V] consensus sequence.

### Statistical analysis

Statistical analysis was performed with Prism 10 software, with specific details about the statistical tests conducted, the number of independent experiments, and P values provided in the corresponding figure legends.

## Acknowledgements

This research was supported by grants from Aligning Science Across Parkinson’s disease (ASAP-000580) through the Michael J. Fox Foundation for Parkinson’s Research (MJFF) and the Ludwig Foundation. Agnes Roczniak-Ferguson (Yale), Devin Clegg (Yale), Timothy Ryan (Weill-Cornell), Kallol Gupta (Yale), Pietro De Camilli (Yale) and Tom Melia (Yale) provided valuable feedback. We appreciate guidance and advice from Wade Harper (Harvard), Annan Cook (UC-Berkeley), Andres Leschziner (UCSD), Jaime Alegrio Louro (UCSD), and Zihao Lin (Yale) with predictions of LRRK2-GABARAP interactions. We thank Berrak Ugur (Yale) for managing our ASAP project. The authors do not have any conflicts to declare.

## Author contributions

ABD and SMF designed experiments. ABD performed all experiments but 7G and S1B. AF performed iPSC-derived macrophage work (S1B) and SF performed experiment 7G. ABD and SMF analyzed data and prepared the manuscript.

## Summary of Supplemental Material

Figure S1 provides additional evidence in support of STING-dependent activation of LRRK2 kinase activity.

Figure S2 demonstrates the morphology of lysosomes treated with DMXAA and LLOME.

Figure S3 contains additional data supporting the CASM-dependent activaiton of LRRK2

Figure S4 further supports that LRRK2 activation is dependent on GABARAP.

Figure S5 demonstrates that CASM-inducing stimuli cause GABARAP accumulation at lysosomes.

Supplemental Tables 1-6 provide information concerning reagents and tools used in this study.

