## Supplemental Material for "A STING-CASM-GABARAP Pathway Activates LRRK2 at Lysosomes"

### Supplemental Figures

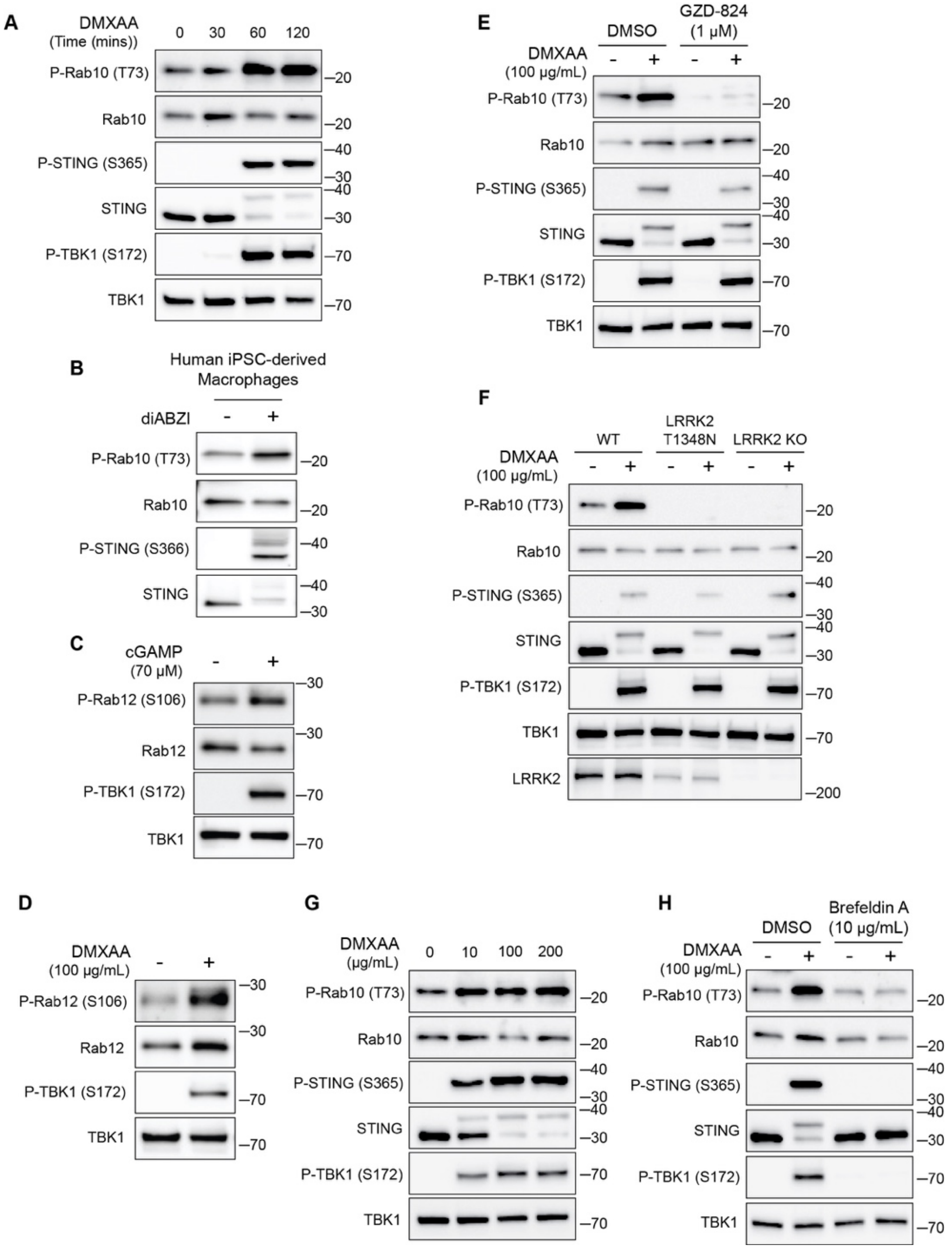

**Figure S1: STING-dependent activation of LRRK2 kinase activity.** (A) Immunoblots illustrating the time course of responses to treatment with 100 μg/ml DMXAA. (B) Immunoblots from human iPSC-derived macrophages treated for 2 hours with 2.5 μM diABZI. (C-D) Immunoblots showing the impact of STING activation for 2 hours with cGAMP and DMXAA on Rab12 phosphorylation (P-TBK1 is a control for STING activation). (E) Immunoblots from cells incubated for 2 hours +/- DMXAA and +/- GZD-824 (LRRK2 inhibitor, treatment started 1 hour before addition of DMXAA). (F) Immunoblots demonstrating the effects of STING activation with DMXAA in WT, LRRK2 T1348N knockin and LRRK2 knockout RAW 264.7 cells. (G) Immunoblots showing the concentration dependence of responses to STING activation with DMXAA. (H) Immunoblots from cells treated with the indicated concentrations of DMXAA (2 hours) and Brefeldin A (treatment started 1 hour before addition of DMXAA). Molecular weight markers (kDa) are indicated on the right side of each blot. All experiments apart from S1D were conducted in RAW 264.7 cells. Data presented in this figure is representative of results from a minimum of 3 independent experiments.

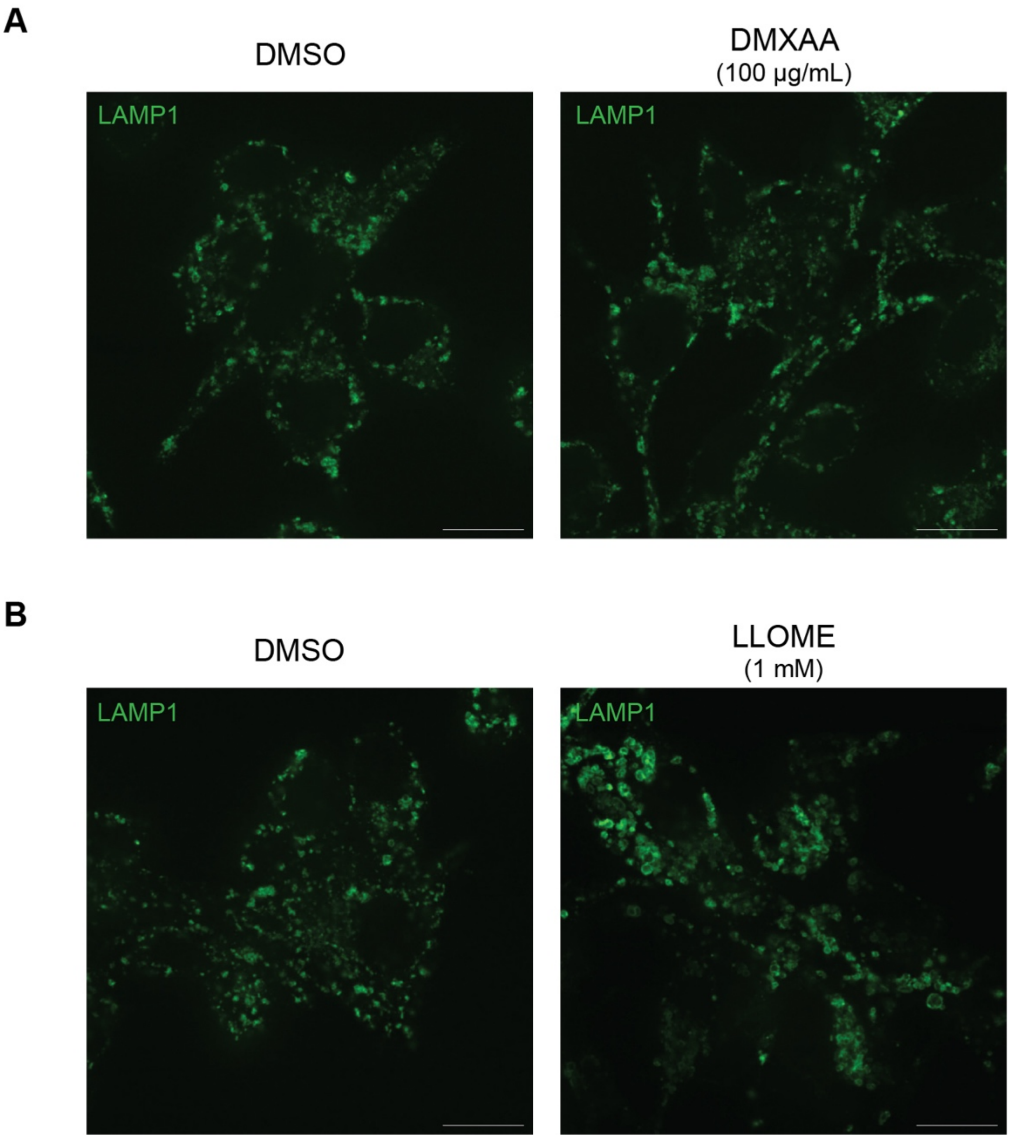

**Figure S2: Morphology of lysosomes treated with DMXAA and LLOME.** (A) Spinning disk confocal immunofluorescence microscopy analysis of lysosome morphology (LAMP1) of WT RAW 264.7 cells treated with the STING agonist DMXAA (100 µg/ml for 2 hours). (B) Spinning disk confocal immunofluorescence microscopy analysis of lysosome morphology (LAMP1) of WT RAW 264.7 cells treated with LLOME (1 mM for 2 hours). The scale bars are 10 µm. Data presented in this figure is representative of results from a minimum of 3 independent experiments.

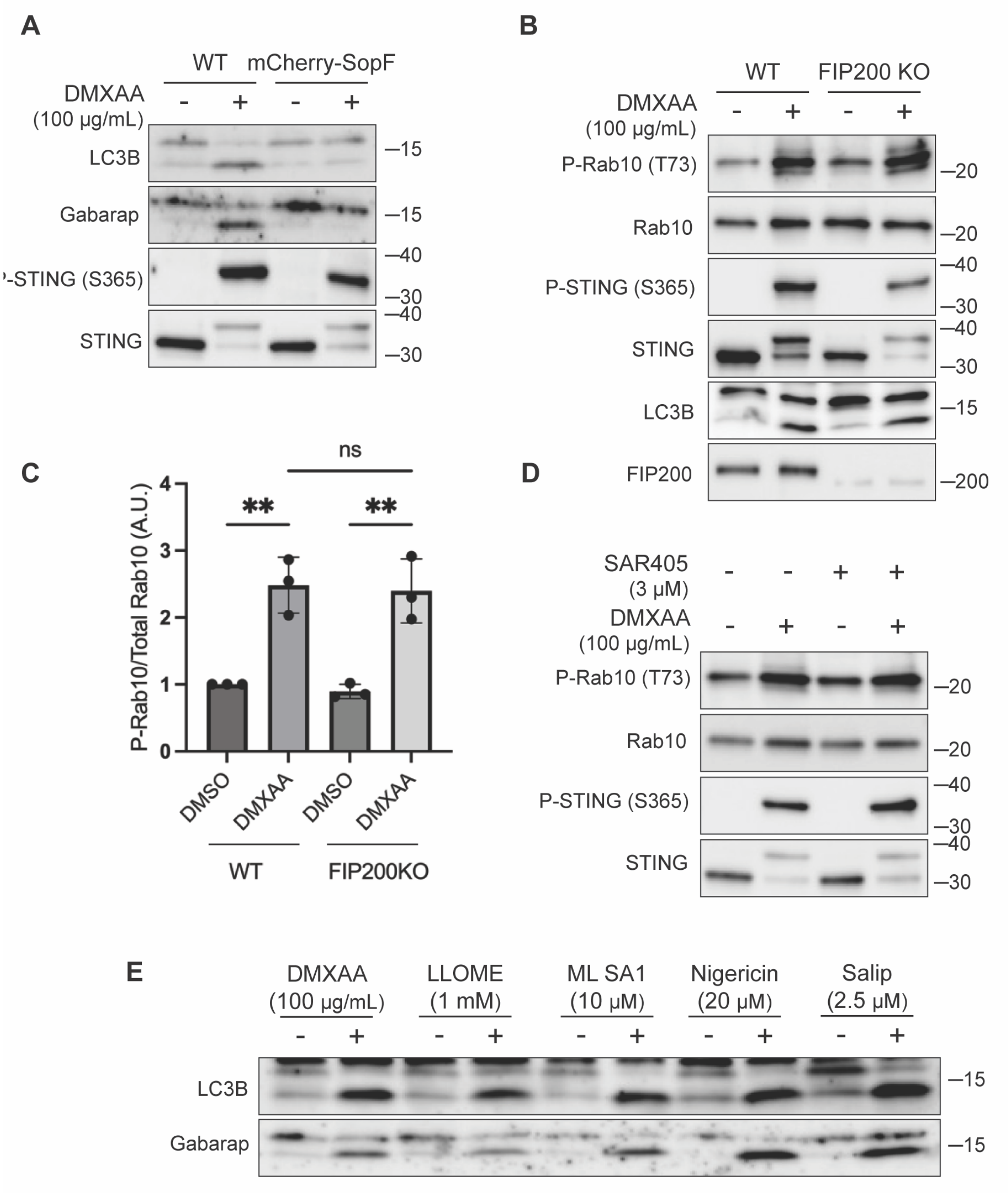

**Figure S3: LRRK2 activation is dependent on CASM.** (A) Immunoblots from WT and mCherry-SopF expressing RAW 264.7 cells treated with DMXAA for 2 hours. (B) Immunoblots from cells treated with DMXAA for 2 hours in FIP200 knockout cells. (C) Quantification of the phospho-Rab10/Rab10 ratios from panel B. A One-Way ANOVA with Sidak's post-test was performed (WT DMSO vs WT DMXAA, P = 0.0029; FIP200 knockout DMSO vs FIP200 knockout DMXAA, P = 0.0027; WT DMXAA vs FIP200 knockout DMXAA, P = 0.9998). Error bars represent standard deviations. (D)

Immunoblots from cells treated with SAR405, a PIK3C3/VPS34 inhibitor, +/- DMXAA. Cells were pretreated +/- SAR405 for 30 minutes prior to the addition of DMXAA and SAR405 was maintained throughout the DMXAA treatment. (E) Immunoblots demonstrating all CASM activating compounds tested cause the lipidation of LC3B and Gabarap after 2 hours of treatment. NEM (20 mM) was used during lysis to help preserve ATG8 lipidation. Molecular weight markers (kDa) are indicated on the right side of each blot. All experiments were conducted in RAW 264.7 cells. Data presented in this figure is representative of results from a minimum of 3 independent experiments.

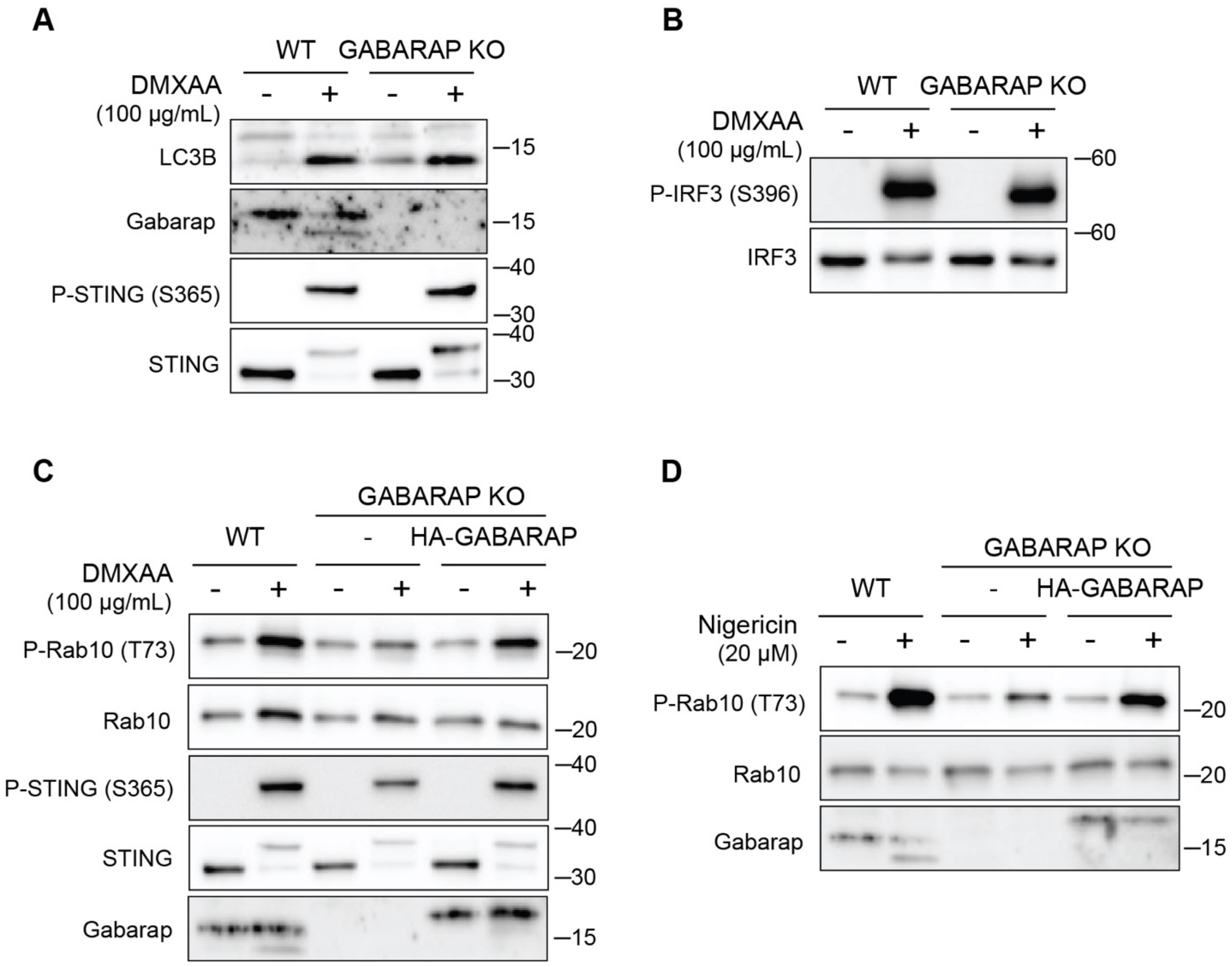

**Figure S4: LRRK2 activation is dependent on GABARAP.** (A) Immunoblots from WT and GABARAP knockout cells treated with DMXAA for 2 hours assessing LC3B and GABARAP lipidation. (B) Immunoblots from WT and GABARAP knockout cells treated with DMXAA for 2 hours to assess IRF3 phosphorylation. (C-D) Immunoblots from WT, GABARAP KO, and GABARAP KO + HA-GABARAP rescue cells treated with DMXAA in C and Nigericin in D for 2 hours. Molecular weight markers (kDa) are indicated on the right side of each blot. All experiments were conducted in RAW 264.7 cells. Data is representative of results from a minimum of 3 independent experiments.

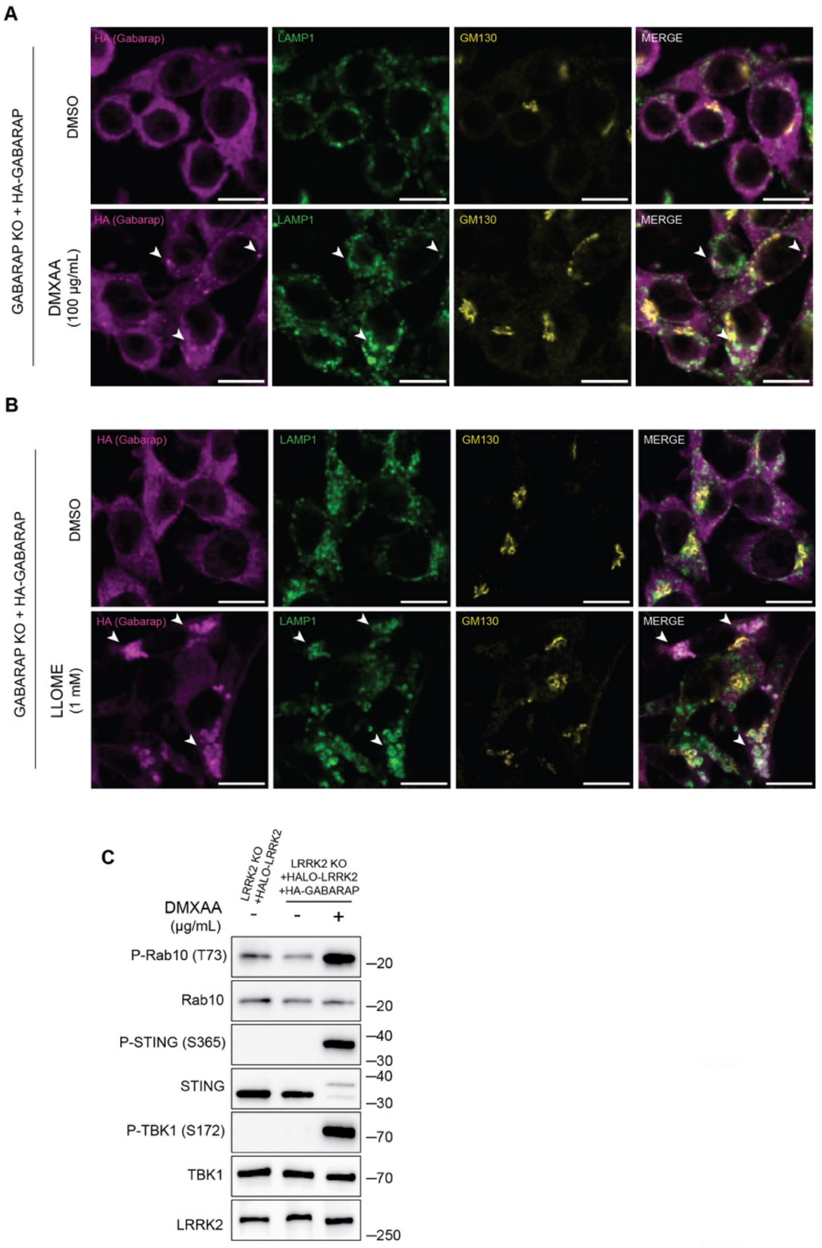

**Figure S5: CASM-inducing stimuli cause GABARAP accumulation at lysosomes.** (A) Immunofluorescence confocal microscopy analysis of HA-GABARAP localization in cells treated with DMXAA for 2 hours. The scale bar represents 10 μm. (B) Immunofluorescence confocal microscopy analysis of HA-GABARAP localization in cells treated with LLOME for 2 hours. The scale bar represents 10 μm. (C) Immunoblots from LRRK2 KO + HALO-LRRK2 + HA-GABARAP cells treated with DMXAA for 2 hours.

Molecular weight markers (kDa) are indicated on the right side of each blot. All experiments were conducted in RAW 264.7 cells. Data is representative of results from a minimum of 3 independent experiments.

### Supplemental Methods

**Table 1: Summary of cell lines used in this study.**

| Cell Line | Genotype | Reference | RRID |
|---|---|---|---|
| RAW 264.7 | WT LRRK2 parental | ATCC SC-6003 | CVCL_UL71 |
| RAW 264.7 | LRRK2 KO | ATCC SC-6004 | CVCL_UL72 |
| RAW 264.7 | LRRK2 T1348N | ATCC SC-6005 | CVCL_UL73 |
| RAW 264.7 | STING KO | Talaia et al 2024 | CVCL_D7F3 |
| RAW 264.7 | STING KO + mouse STING (WT) | This paper | CVCL_D7F4 |
| RAW 264.7 | STING KO + mouse STING (amino acids 1-339) | This paper | CVCL_D7F5 |
| RAW 264.7 | TBK1 KO | This paper | CVCL_D7F6 |
| RAW 264.7 | IKKε KO | This paper | CVCL_D7F1 |
| RAW 264.7 | TBK1 KO + IKKε KO | Talaia et al 2024 | CVCL_D7F7 |
| RAW 264.7 | Atg16L1 KO | This paper | CVCL_D7EZ |
| RAW 264.7 | FIP200 KO | This paper | CVCL_E2WW |
| RAW 264.7 | mCherry-SopF | This paper | CVCL_D7F2 |
| RAW 264.7 | GABARAP KO | This paper | CVCL_D7F0 |
| RAW 264.7 | GABARAP KO + HA-human GABARAP | This paper | CVCL_E2WX |
| RAW 264.7 | LRRK2 KO + HALO-human LRRK2 | This paper | CVCL_D7F8 |
| RAW 264.7 | LRRK2 KO + HA-human GABARAP | This paper | CVCL_E2WY |
| RAW 264.7 | LRRK2 KO + HALO-human LRRK2 + HA-human GABARAP | This paper | CVCL_D7F9 |
| RAW 264.7 | LRRK2 KO + HALO-human LRRK2 (LIR1 mutant) + HA-human GABARAP | This paper | CVCL_E2WZ |
| RAW 264.7 | LRRK2 KO + HALO-human LRRK2 (LIR2 mutant) + HA-human GABARAP | This paper | CVCL_E2X0 |

| | | | |
|---|---|---|---|
| RAW 264.7 | LRRK2 KO + HALO-human LRRK2 (LIR1+2 mutant) + HA-human GABARAP | This paper | CVCL_E2X1 |
| IPSC A18945 | | | CVCL_RM92 |

#### Table 2: Summary of supplies for cell culture, drug treatments and lab chemicals

| **Cell culture reagents** | | |
|---|---|---|
| **Reagent** | **Company** | **Product Number** |
| DMEM | Thermo Fisher Scientific | 11965-092 |
| E8 | Thermo Fisher Scientific | A15169-01 |
| E8 Supplement | Thermo Fisher Scientific | A15171-01 |
| RPMI | Thermo Fisher Scientific | 11875093 |
| Rock inhibitor | Stemcell Technologies | 100-1044 |
| STEMdiff Hematopoietic Kit | Stemcell Technologies | 5310 |
| Matrigel | Corning | 356230 |
| Recombinant Human M-CSF | Peprotech | 300-25 |
| FBS | Thermo Fisher Scientific | 16140-071 |
| PBS | Thermo Fisher Scientific | 10010023 |
| Cell Stripper | Corning | 25056CI |
| Penicillin/Streptomicin (10,000 U/mL) | Thermo Fisher Scientific | 15140122 |
| Blasticidin | Invivogen | Ant-bl-05 |
| Puromycin | Thermo Fisher Scientific | A11138-03 |
| Opti-Mem | Thermo Fisher Scientific | 31985062 |
| Lipofectamine 2000 | Invitrogen | 11668019 |
| Lipofectamine RNAiMAX | Invitrogen | 2448190 |
| Lipofectamine CRISPRMAX | Invitrogen | CMAX00003 |
| Fugene HD | Promega | E2311 |
| **Drugs/Compounds** | | |
| **Compound** | **Company** | **Product Number** |
| 2,3 cGAMP | Chemietek | CT-CGMAP |
| DMXAA | Cayman Chemicals | 14617 |
| diABZI | Cayman Chemicals | 28054 |
| MLi-2 | Abcam | Ab254528 |
| GZD-824 | Cayman Chemicals | 21508 |
| SAR405 | Selleck Chemicals | S7682 |
| Nigericin | Cayman Chemicals | 11437 |
| ML-SA1 | Cayman Chemicals | 29958 |
| Saliphenylhalamide | Omm Scientific | N/A |

| | | |
|---|---|---|
| Folimycin | Abcam | Ab144277 |
| N-Ethylmaleimide | Sigma-Aldrich | E3876 |
| **Lab Supplies** | | |
| **Reagent** | **Company** | **Product Number** |
| Potassium Phosphate Monobasic | J.T. Baker | 3246-01 |
| Sodium Phosphate Dibasic | J.T. Baker | 3828-05 |
| Glycine | American Bio | AB00730-05000 |
| Tris | American Bio | AB02000-05000 |
| NaCl | Sigma-Aldrich | 3624-05 |
| Hydrochloric Acid | J.T. Baker | 9535 |
| SDS | American Bio | AB01920-00500 |
| EDTA | Sigma-Aldrich | 03690 |
| Triton X-100 | Sigma-Aldrich | X100 |
| Tween-20 | Sigma-Aldrich | P7949 |
| Glycerol | American Bio | AB00751 |
| Bromphenol Blue | Sigma-Aldrich | B5525 |
| B-mercaptoethanol | Sigma-Aldrich | M3148 |
| Sucrose | Sigma-Aldrich | S0389 |
| EGTA | Sigma-Aldrich | E4378 |
| HEPES (pH 7.4) | Thermo Fisher Scientific | 15630-080 |
| DMSO | Sigma-Aldrich | D2650 |
| COmplete mini EDTA Free | Roche | 11836170001 |
| PhosSTOP | Roche | 4906837001 |
| Coomassie Plus Protein Assay Reagent | Thermo Fisher Scientific | 23236 |
| PAGEruler Plus Prestained Protein Ladder | Thermo Fisher Scientific | 26620 |
| Biotin Protein Ladder | Cell Signaling | 7727L |
| 4-15% MiniPROTEAN 10-well | Biorad | 4568084g |
| 4-15% MiniPROTEAN 12-well | Biorad | 4568085 |
| 4-15% MiniPROTEAN 15-well | Biorad | 4568086 |
| BSA | Sigma-Aldrich | A9647 |
| Non-Fat Dry Milk Omniblock | American Bio | AB10109-01000 |
| 0.45 um Nitrocellulose Membrane | Thermo Fisher Scientific | 1620115 |
| Whatman Filter Paper | VWR | 28298-020 |
| SuperSignal West Pico PLUS Chemiluminescence Substrate | Thermo Fisher Scientific | 34580 |
| SuperSignal West Femto Maximum Sensitivity Substrate | Thermo Fisher Scientific | 34095 |
| Methanol | Sigma-Aldrich | 179337-4L-PB |
| Ethanol | Decon Laboratories | 2716 |
| Ampicillin | Sigma-Aldrich | A0166 |

| | | |
|---|---|---|
| Tryptone | RPI | T600-60 |
| LB + Ampicillin (100 µg/mL) | Recombinant Technologies | 760100 |
| Iron (II) Chloride | Sigma-Aldrich | 220299 |
| Iron (III) Chloride | Sigma-Aldrich | 157740 |
| Ammonium hydroxide (30%) | Sigma-Aldrich | 320145 |
| Dextran | Sigma-Aldrich | D1662 |
| Snakeskin dialysis tubing (10,000 Mol Wt) | Thermo Fisher Scientific | 68100, 10,000 |
| LS Columns | Miltenyi Biotec | 130-042-401 |
| QuadroMACS Separator | Miltenyi Biotec | 130-091-051 |
| Pierce Anti-HA Magnetics Beads | Thermo Fisher Scientific | 88837 |
| Saponin Quilajja sp. | Sigma-Aldrich | S4521 |
| Paraformaldehyde | Electron Microscopy Sciences | 19202 |
| Sodium dihydrogen phosphate monohydrate | J.T. Baker | 3818 |
| Sodium phosphate, dibasic, anhydrous | J.T. Baker | 3828 |
| ProLong™ Gold Antifade Mountant with DNA Stain DAPI | Thermo Fisher Scientific | P36935 |
| Fisherbrand™ Superfrost™ Disposable Microscope Slides | Thermo Fisher Scientific | 12-550-143 |
| Microscope Cover Slips (12 mm) | Carolina Biological Supply | 633029 |
| **Molecular Biology Reagents** | | |
| **Reagent** | **Company** | **Product Number** |
| Q5 High-Fidelity 2X Master Mix | NEB | M0492S |
| HIFI DNA Assembly Master Mix | NEB | E2621L |
| One-Shot STABL3 | Invitrogen | C7373-03 |
| **Software** | | |
| **Version** | **Company** | **RRID** |
| Prism 10 | Graphpad | SCR_002798 |
| 2.14.0/1.54f | FIJI | SCR_002285 |
| 1.7.1 | ChimeraX | |
| AlphaFold 2.2.4 | Google Deepmind | |
| AlphaFold Server | Google Deepmind | |
| ChatGPT-4o | OpenAI | |

### Table 3: Summary of plasmids used in this study

| Plasmid | Reference | RRID |
|---|---|---|
| pPB-EF1A-HALO-hLRRK2 | This paper | In Progress |
| pPB-EF1A-mSTING | This paper | In Progress |

| | | |
|---|---|---|
| pPB-EF1A-mSTING (1-339) | This paper | In Progress |
| pPB-HA-hGABARAP | This paper | In Progress |
| pPB-EF1A-HALO-hLRRK2 (LIR1 mutant WEVL -> AEVA) | This paper | In Progress |
| pPB-EF1A-HALO-hLRRK2 (LIR2 mutant WTFI -> ATFA) | This paper | In Progress |
| pPB-EF1A-HALO-hLRRK2 (LIR1+2 mutant WEVL -> AEVA / WTFI -> ATFA) | This paper | In Progress |
| mCherry-SopF | Addgene (135174) | Addgene_135174 |
| pPB-mCherry-SopF | This paper | In Progress |
| pEIF1a-Piggybac transposase | Michael Ward (NINDS)(Pantazis et al., 2022) | |

### Table 4: Sequences of oligonucleotide primers used in this study

| Primer | Primer | Reference |
|---|---|---|
| mSTING (1-339)_F | TGAACCCAGCTTTCTTGTAC | This paper |
| mSTING (1-339)_R | CTCCTCCTTTTCTTCCTG | This paper |
| MCherry-SopF Ins_F | tccatttcaggtgtcgtgacGGTTTAGTGAACCGTCAG | This paper |
| MCherry-SopF Ins_R | tttgtacaagaaagctgggtTCAATATAATATTATGCAGTCTCTATTAAG | This paper |

### Table 5: Description of antibodies used in this study

| **Antibodies** | | | | |
|---|---|---|---|---|
| **Antibody** | **Company** | **Product Number** | **Concentration** | **RRID** |
| LRRK2 | Abcam | Ab133474 | 1:2000 | AB_2713963 |
| STING | Cell Signaling Technologies | 13647S | 1:1000 | AB_2732796 |
| P-STING S365 | Cell Signaling Technologies | 72971S | 1:1000 | AB_2799831 |
| P-STING S366 | Cell Signaling Technologies | 19781S | 1:1000 | AB_2737062 |
| TBK1 | Cell Signaling Technologies | 3504S | 1:2000 | AB_2255663 |
| P-TBK1 S172 | Cell Signaling Technologies | 5483S | 1:2000 | AB_10693472 |
| IKKε | Cell Signaling Technologies | 2905S | 1:2000 | AB_1147662 |
| Rab10 | Cell Signaling Technologies | 8127S | 1:1000 | AB_10828219 |
| P-Rab10 T73 | Abcam | Ab241060 | 1:1000 | AB_2884876 |
| Rab12 | Santa Cruz | Sc-515613 | 1:500 | AB_3101762 |
| P-Rab12 S106 | Abcam | Ab256487 | 1:1000 | AB_2884880 |

| | | | | |
|---|---|---|---|---|
| Atg16L1 | Cell Signaling Technologies | 8089S | 1:2000 | AB_10950320 |
| LC3B | Thermo Fisher Scientific | PA1-46286 | 1:1000 | AB_2234770 |
| GABARAP | Cell Signaling Technologies | 13733S | 1:1000 | AB_2798306 |
| FIP200 | Cell Signaling Technologies | 12436S | 1:2000 | AB_2797913 |
| LAMP1 (1D4B) | DSHB | AB-528127 | 1:12000<br>1:400 | AB_2134500 |
| PDI | Cell Signaling Technologies | 2446S | 1:1000 | AB_2298935 |
| GM130 | BD Biosciences | 610822 | 1:1000<br>1:200 | AB_398141 |
| HA | Sigma-Aldrich/Roche | 12013819001 | 1:1000 | AB_390917 |
| HA | Cell Signaling Technologies | 3724S | 1:100 | AB_1549585 |
| Rabbit IgG (HRP) | Cell Signaling Technologies | 7074S | 1:2000 | AB_2099233 |
| Mouse IgG (HRP) | Cell Signaling Technologies | 7076S | 1:2000 | AB_330924 |
| Rat IgG (HRP) | Cell Signaling Technologies | 7077S | 1:2000 | AB_10694715 |
| Biotin (HRP) | Cell Signaling Technologies | 7075S | 1:4000 | AB_10696897 |
| AlexaFluor 488 anti-rat | Invitrogen | A21208 | 1:600 | AB_2535794 |
| AlexaFluor 568 anti-mouse | Invitrogen | A10037 | 1:600 | AB_11180865 |
| AlexaFluor 647 anti-rabbit | Invitrogen | A31573 | 1:600 | AB_2536183 |

### Table S6: Summary of siRNAs used in this study

| **siRNA** | **Horizon Biosciences Product Number** |
|---|---|
| Negative Control | D-001206-13-05 |
| Lrrk2 | M-049666-01-0005 |
| Rab10 | M-040862-01-0005 |
| Rab12 | M-040865-01-0005 |
| Atg3 | M-048439-02-0005 |
| Atg16L1 | M-051699-01-0005 |
| Map1lc3a | M-056203-00-0005 |
| Map1lc3b | M-040989-01-0005 |
| Gabarap | M-041776-01-0005 |
| GabarapL1 | M-040444-01-0005 |
| GabarapL2 | M-059605-01-0005 |
